# PGR5 is needed for redox-dependent regulation of ATP synthase both in chloroplasts and in cyanobacteria

**DOI:** 10.1101/2024.11.03.621747

**Authors:** Lauri Nikkanen, Laura T. Wey, Russell Woodford, Henna Mustila, Darius Kosmützky, Maria Ermakova, Eevi Rintamäki, Yagut Allahverdiyeva

**Affiliations:** Molecular Plant Biology, Department of Life Technologies, University of Turku, Finland; School of Biological Sciences, Monash University, Australia; Department of Biochemistry, University of Cambridge, United Kingdom

**Author notes:** Co-corresponding authors **Authors for Correspondence:** Lauri Nikkanen, Molecular Plant Biology, Department of Life Technologies, University of Turku, Turku 11 FI-20014, Finland. Yagut Allahverdiyeva, Molecular Plant Biology, Department of Life Technologies, University of Turku, Turku 11 FI-20014, Finland.

**Keywords:** Photosynthesis, ATP synthase, proton motive force, chloroplast, cyanobacteria, Arabidopsis, electrochromic shift

## Abstract

The regulation of proton motive force (*pmf*) via ATP synthase activity is a critical mechanism by which photosynthetic organisms maintain redox homeostasis and activate photoprotective responses under fluctuating light conditions. Here, we used time-resolved electrochromic shift (ECS) measurements to investigate *pmf* dynamics across a range of photosynthetic organisms. Our results reveal that ATP synthase is dynamically regulated during light fluctuations, but in *Arabidopsis thaliana* could this regulation could not be explained by the known light-induced reduction of the CF□γ subunit, suggesting alternative control mechanisms. The PROTON GRADIENT REGULATION 5 (PGR5) protein, previously proposed to facilitate cyclic electron transport (CET), also has a potential role in ATP synthase regulation. The physiological role of cyanobacterial Pgr5 has remained elusive. We characterised a Δ*pgr5* mutant of *Synechocystis* sp. PCC 6803 and investigated *pmf* dynamics in *pgr5* mutants of *Chlamydomonas reinhardtii*, Arabidopsis, and the C4 grass *Setaria viridis*. While PGR5 was not required for CET in *Synechocystis*, it was needed for downregulating ATP synthase under high irradiance in all tested organisms via a thiol redox state dependent mechanism. Furthermore, interaction between AtPGR5 and AtCF□γ suggests that PGR5 functions as a conserved inhibitor of ATP synthase under high light, contributing to *pmf* retention and photoprotection.

**Highlight:** PGR5 is involved in regulation of ATP synthase in fluctuating light conditions across photosynthetic organisms in a manner that depends on the thiol redox state of the chloroplast or cell.

## INTRODUCTION

Oxygenic photosynthesis by plants, algae, and cyanobacteria uses light energy to drive oxidation of water to produce ATP and NADPH for CO_2_ fixation and other cellular processes. In their natural habitats, oxygenic photosynthetic organisms undergo constant fluctuation of irradiance. Mechanisms to adjust the redox homeostasis in chloroplasts and cyanobacterial cells have evolved to prevent energy-consuming futile reactions and damage to the photosynthetic machinery and other biomolecules via generation of reactive oxygen species (ROS) by the photosynthetic apparatus. ROS can be generated as a result of excitation imbalance between the photosystems, or when electron transport rate exceeds the sink capacity of the reactions downstream of the photosynthetic electron transport chain (PETC). Resulting accumulation of reduced electron carriers in the PETC triggers charge recombination reactions within the photosystems and direct reduction of O_2_, generating highly reactive ROS such as superoxide (O_2_^•−^) and hydrogen peroxide (H_2_O_2_) at PSI and singlet oxygen (^1^O_2_) primarily at PSII (Foyer & Hanke, 2022). PSI and its iron sulphur clusters are particularly vulnerable to over-reduction-induced damage, and plants, algae, and cyanobacteria invest heavily in their protection (Tiwari et al., 2016, 2024). These protective mechanisms involve cyclic electron transport around PSI (CET), redistribution of excitation energy between the photosystems via state transitions, non-photochemical quenching (NPQ), and inhibition of PQH_2_ oxidation at the cytochrome b_6_f complex, known as photosynthetic control (Tikkanen & Aro, 2014; Shimakawa et al., 2016; Nikkanen et al., 2021; Eckardt et al., 2024; Degen & Johnson, 2024). Additionally, in photosynthetic organisms apart from angiosperms, flavodiiron proteins provide a photoprotective electron sink on the acceptor side of PSI by catalysing controlled reduction of O_2_ to water (the Mehler-like reaction) (Allahverdiyeva et al., 2013; Ilik et al., 2017).

As photosynthetic control and NPQ (in plants and algae) are induced by acidic pH of the thylakoid lumen, an important mechanism to maintain photosynthetic redox balance involves control of the proton motive force *(pmf)* via regulation of the thylakoid ATP synthase (Kanazawa et al., 2017; Takagi et al., 2017). The pH-dependent component of *pmf* (ΔpH), is generated by release of protons to the lumenal side of the thylakoid membrane by water oxidation at PSII and the Q cycle, and proton pumping from the stromal/cytosolic side by cyclic electron transport, as well as consumption of protons on the stromal side. The proton gradient is released mainly via efflux through the ATP synthase, providing the driving force for ATP synthesis. Proton-conductivity of the ATP synthase is therefore an important regulatory factor for adjusting the magnitude of the *pmf*.

In addition to ΔpH, the *pmf* comprises an electric field component (ΔΨ). ΔΨ derives from charge separation within the photosystems and in the Q cycle, the positive charge of protons, and exchange of cations (e.g. K^+^, Ca^2+^ and Mg^2+^) in and anions (e.g. Cl^-^) out of the lumen by ion channels and transporters in the thylakoid membrane (Armbruster et al., 2017). ΔpH and Δψ are thermodynamically equivalent and both drive ATP synthesis (Hangarter & Good, 1982). However, build-up of sufficient ΔpH over the thylakoid membrane is required for induction of photoprotective mechanisms upon sudden increases in light intensity, namely dissipation of excessive excitation energy as heat in NPQ (Niyogi & Truong, 2013; Krishnan-Schmieden et al., 2021) and photosynthetic control (Tikhonov et al., 1981; Degen & Johnson, 2024). These mechanisms prevent over-reduction of the electron transport chain and photodamage to PSI. Conversely, it is also important to down-regulate ΔpH (and consequently, NPQ and photosynthetic control) upon sudden decreases in irradiance in order to maintain sufficient supply of electrons on the acceptor side of the electron transfer chain for carbon fixation in the Calvin–Benson cycle (CBC).

ATP synthase is regulated via several mechanisms. The F_1_ε (epsilon) subunits of the chloroplast (CF_0_CF_1_) ATP synthase (Feniouk et al., 2006) and the cyanobacterial ATP synthase (Imashimizu et al., 2011; Murakami et al., 2018) have inhibitory roles preventing ATP hydrolysis. F_1_ε undergoes conformational changes triggered by *pmf* generation and changes in ATP/ADP ratio, allowing inhibition of ATP hydrolysis (Richter & McCarty, 1987; Feniouk et al., 2006). Moreover, upon dark-to-light transitions, the chloroplast ATP synthase is activated by thioredoxin (TRX)-mediated reduction of a disulphide in the γ subunit (CF_1_γ) (McKinney et al., 1978; Nalin & McCarty, 1984; Akiyama et al., 2023). CF_1_γ is kept in an inactive oxidised state in darkness to avoid hydrolysis of ATP by reverse activity or loss of the proton gradient (Hisabori et al., 2013; Kohzuma et al., 2017; Hahn et al., 2018). Interestingly, it was recently reported that the *Chlamydomonas reinhardtii* CF_1_γ, although the conserved disulphide is present, contains a modified hairpin loop in the redox regulatory domain that allows CrCF_0_CF_1_ to remain active also in darkness to facilitate heterotrophic metabolism (Lebok & Buchert, 2024). In plants, maintenance of adequate dark-*pmf* via ATP synthase dark-light regulation is also needed for powering the insertion of new proteins into the thylakoid membrane via the Tat- and Sec-pathways to maintain the integrity of the photosynthetic apparatus (Kohzuma et al., 2017). CF_1_γ is reduced in seconds upon illumination, which lowers the threshold *pmf* for activation of the ATP synthase (Kramer et al., 1990; Konno et al., 2012; Kohzuma et al., 2017; Nikkanen et al., 2018; Buchert et al., 2021). In turn, ΔpH also facilitates reduction of the CF_1_γ disulphide bond (Sekiguchi et al., 2024), while dissipation of ΔpH promotes oxidation of CF_1_γ (Sekiguchi et al., 2022). F- and m-type chloroplast TRX isoforms have been understood to be mainly responsible for light-dependent reduction of CF_1_γ (Schwarz et al., 1997; Sekiguchi et al., 2020), and chloroplast TRX-like proteins (TrxL2) and 2-cysteine peroxiredoxins (2-CysPrxs) catalyse the oxidation of the subunit in the dark (Yoshida et al., 2018; Sekiguchi et al., 2022). Additionally, *in vivo* studies have revealed the involvement of the NADPH-dependent TRX system (NTRC) in CF_1_γ reduction in low light conditions (Carrillo et al., 2016; Nikkanen et al., 2016), while the atypical TRX CDSP32 was recently suggested to play a role in reduction of CF_1_γ in potato (Rey et al., 2024).

Reduction of CF_1_γ constitutes an on/off switch controlling the activation of the ATP synthase upon the onset of illumination in chloroplasts of plants and algae, and does not occur in mitochondria or in cyanobacteria (Hisabori et al., 2013). The cyanobacterial γ subunit still has a role in preventing ATP hydrolysis however, via an ADP-dependent mechanism (Sunamura et al., 2010). Moreover, the small AtpΘ protein (Song et al., 2022) has been reported to function as an inhibitor of the cyanobacterial ATP synthase in the dark and in low *pmf* conditions, preventing ATP hydrolysis and collapse of the proton gradient. Interestingly, the F_1_α (AtpA) and F_1_β (AtpB) subunits of the ATP synthase also contain conserved Cys residues, and F_1_α has been identified as a prospective TRX target in cyanobacteria (Lindahl & Florencio, 2003; Guo et al., 2014), and F_1_β in plants (Balmer et al., 2006). Moreover, the mitochondrial ATP synthase in *Xenopus laeviis* oocytes is regulated by reversible oxidation of thiols in the F_1_α subunit (Cobley et al., 2020).

There are also other regulatory mechanisms that adjust the activation state of the ATP synthase according to the metabolic state of the stroma or cytosol, such as substrate (ADP and P_i_) availability; a lack of Pi due to low consumption of ATP by an inactive CBC would lead to a decrease ATP synthase activity, increasing ΔpH and activating photosynthetic control and NPQ to protect PSI from over-reduction (Kanazawa & Kramer, 2002; Avenson et al., 2005; Takagi et al., 2017). High CO_2_ supply results in increase of chloroplast ATP synthase activity by a mechanism that is independent of CF_1_γ redox state (Kiirats et al., 2009; Kohzuma et al., 2013). Another potential regulatory mechanism has been suggested to involve phosphorylation of the CF_1_β subunit (del Riego et al., 2006), although a more recent study reports that the phosphorylation is unlikely to have a role in controlling ATP synthase activity and is instead involved in CF_1_ assembly (Strand et al., 2023).

Additionally, thylakoid conductivity and the *pmf* can be adjusted through the activities of ion channels and transporters in the thylakoid membrane (Spetea et al., 2017). A lack of the K^+^/H^+^ exchanger KEA3 or the Cl^-^ channels CLCe and VCCN1 has been shown to affect the *pmf* and/or induction and relaxation of NPQ in plants (Armbruster et al., 2014; Herdean et al., 2016b,a; Wang & Shikanai, 2019; von Bismarck et al., 2023). Similarly, lack of the cyanobacterial K^+^ channel SynK resulted in decreased ΔpH generation and impairment of photosynthetic control in the model cyanobacterium *Synechocystis* sp. PCC 6803 (*Synechocystis*) (Checchetto et al., 2012). Controlling the ionic environment of the thylakoid membrane is particularly important to avoid drastic increases in ΔΨ, which result in production of damaging ^1^O_2_ via recombination reactions in PSII (Davis et al., 2016).

Plants deficient in the PROTON GRADIENT REGULATION 5 (PGR5) protein suffer from impaired *pmf* generation (Munekage et al., 2002). In plant chloroplasts, PGR5 forms redox-dependent heterodimers with PGRL1 (PGR5-LIKE 1), which have been proposed to mediate or regulate cyclic electron transport (CET) from ferredoxin to the plastoquinone pool (DalCorso et al., 2008; Hertle et al., 2013; Yamamoto & Shikanai, 2019), contributing to *pmf* formation (Wang et al., 2015). More recently, a functional PGRL1-analogue was identified also in cyanobacteria (Dann & Leister, 2019).

The *pgr5* knockout plants are unable to induce photosynthetic control and NPQ during transitions from low to high light (HL) (Munekage et al., 2002; Suorsa et al., 2012; Yamamoto & Shikanai, 2019). Consequently, exposure of *pgr5* mutant plants to HL results in over-reduction of the electron transport chain and photodamage to the iron sulphur clusters of PSI (Tiwari et al., 2016, 2024).

Accordingly, the lack of PGR5 is lethal in fluctuating light conditions in *Arabidopsis thaliana (Arabidopsis)* (Tikkanen et al., 2010; Suorsa et al., 2012) and severely inhibits growth of the model C_4_ plant *Setaria viridis* (Woodford et al., 2024) as well as of the model green alga *Chlamydomonas reinhardtii (Chlamydomonas)* (Jokel et al., 2018). *Synechocystis* cells deficient in a PGR5-orthologue (SynPgr5) suffer from slightly increased HL-sensitivity (Yeremenko et al., 2005), but no growth impairment was observed in fluctuating light conditions (Allahverdiyeva et al., 2013). Interestingly, however, there is strong selection pressure against overexpression of SynPgr5 (Margulis et al., 2020).

The physiological functions of PGR5 and PGRL1 remain controversial, with some reports linking the proteins to regulation of linear instead of cyclic electron transport from PSI (Nandha et al., 2007; Joliot & Johnson, 2011; Johnson et al., 2014; Tikkanen et al., 2015; Takagi & Miyake, 2018; Rantala et al., 2020; Maekawa et al., 2024), or regulation of the Q-cycle at the cytochrome b_6_*f* complex (Buchert et al., 2020). Impaired *pmf* generation and induction of photoprotection in the *pgr5* mutant could also derive from the inability of the mutant plants to limit the conductivity of the thylakoid membrane under HL (Avenson et al., 2005; Nikkanen et al., 2018; Degen et al., 2023). Interestingly, elevated thylakoid (ATP synthase) conductivity upon HL exposure was not observed in isolated and ruptured *pgr5* mutant chloroplasts (Wang et al., 2018). This suggests that a stromal component that is lost in ruptured chloroplasts plays a role in the effect of PGR5 on thylakoid conductivity.

In the current study, we elucidated the dynamics of *pmf* regulation under light in a variety of photosynthetic organisms focusing on the potential role of PGR5. We performed *in vivo* characterisation of photosynthetic electron transport and *pmf* formation in *Synechocystis, Chlamydonomas*, *Arabidopsis,* and *S. viridis.* Our results reveal that in *Arabidopsis*, dynamic regulation of ATP synthase conductivity during fluctuations in light intensity is independent of the thiol redox state of the CF_1_γ subunit. We show that downregulation of the ATP synthase activity upon transitions to HL depends on PGR5 as well as the thiol redox state of the stroma or cytosol in both chloroplasts and cyanobacteria. Unlike in chloroplasts, PGR5 is not essential for oxidation of the PETC in cyanobacteria. Moreover, we demonstrate that cyanobacterial SynPgr5 is not needed for CET and suggest that the observed effect of plant PGR5 on cyclic electron transport may be an indirect consequence of ATP synthase misregulation and increased photoinhibition of PSI.

## MATERIALS AND METHODS

### Strains, plant lines and growth conditions

*Arabidopsis thaliana* WT-gl1 or WT-Col-0, *pgr5-1* (At2g05620 knockout) (Munekage et al., 2002), and *pgr5-Cas* (At2g05620 CRISPR-Cas9 mutant line) (Penzler et al., 2022) plants were grown for 4 weeks under 200 µmol photons in a 12/12h photoperiod before collecting individual leaves for experiments.

For the BiFC experiments, WT *Nicotiana benthamiana* plants were grown under 130 μmol photons m^−2^Js^−1^ at 23°C in a 16/8 hr light/dark photoperiod.

*Chlamydomonas reinhardtii* WT 137c (progenitor strain of the CC-124 and CC-125 WT strains), *pgr5* knockout and PGR5C complementation strains (Johnson et al., 2014; Jokel et al., 2018) were grown photoautotrophically under continuous 50 μmol photons m^-2^ s^-1^ at 25 °C and 120 rpm in TP medium for four days.

*Setaria viridis cv* MEO V34-1 *pgr5* knock-out mutant was generated using CRISPR-Cas9 gene-editing as described in detail in (Ermakova et al., 2024). WT plants and plants with *pgr5-3* null alleles described in (Woodford et al., 2024) were analysed. Plants were grown in a controlled environment room with a 16□h : 8□h, light : dark photoperiod, 28°C day and 22°C night, 60% humidity and ambient CO_2_ at a light intensity of 300□μmol□m^−2^□s^−1^ supplied by 1000□W red sunrise 3200K lamps (Sunmaster Growlamps, Solon, OH, USA). Youngest fully expanded leaves of 3–4-week-old plants were used in analysis.

*WT,* Δ*pgr5,* Δ*flv3* (Helman et al., 2003) and Δ*pgr5 flv3 Synechocystis* sp. PCC 6803 strains as well as WT *Synechococcus elongatus* sp. PCC 7942 cells were grown under continuous 50 μmol photons m^-2^ s^-1^ at 30 °C and air-level [CO_2_] in pH 7.5 BG-11 medium. Unless stated otherwise, cultures were grown for 4 days in 30 ml flasks, then pelleted and resuspended in fresh BG-11 at an appropriate chlorophyll concentration for each experiment. For the growth curves under fluctuating light, WT, Δ*pgr5* and Δ*pgr5 flv3* cells were grown for 7 days in conditions where light intensity was held at 20 µmol photons m^-2^s^-1^ and increased to 500 µmol photons m^-2^s^-1^ for 30 sec every 5 min. Samples were taken and OD_750 nm_ measured with a spectrophotometer every 24h. For the experiment in Figure 5, cells were grown in a regime in which light intensity fluctuated between 1 min periods of 25 and 500 µmol photons m^-2^s^-1^.

### Construction of Δ*pgr5* and Δ*pgr5 flv3* knockout mutants

The upstream and downstream flanking sequences of the *pgr5* (*ssr2016*) gene and the *aphI* (Km^R^) gene replacing the *ssr2016* open reading frame were amplified from genomic DNA of Δ*pgr5* mutant generated earlier in a different *Synechocystis* WT background as described in (Allahverdiyeva et al., 2013). After transformation into *Synechocystis* sp. PCC 6803 WT strain and the Δ*flv3* strain (Helman et al., 2003), the transformants were selected on BG-11 plates with 50 μg/ml kanamycin. Complete segregation of the mutation was confirmed by PCR with specific forward primer 5′-AGCCAAACCGCATCTAACG-3′ and reverse primer 5′-TTCATAACGCCTCCGGTC-3′. The expected PCR fragment length was 0.7 kb for the WT and 1.7 kb for the mutant (Supplementary Fig. S1). The mutation in the *flv3* open reading frame was confirmed by using forward primer 5′-GTACGGCATGTTCACTACC-3′ and reverse primer 5′-GATCAGCGTTAGCCTGTAA-3′, and the expected PCR fragment length was 1.0 kb for the WT and 2.6 kb for the mutant (Supplementary Fig. S1). Two separate clones of Δ*pgr5* were used interchangeably for the experiments, as no difference was detected in their phenotypes.

### Electrochromic shift (ECS) measurements

Individual *Arabidopsis* leaves were measured with the Dual-PAM 100 spectrophotometer (Walz) and its P515/535 accessory module (Schreiber & Klughammer, 2008) as described earlier (Nikkanen et al., 2018; Guinea Diaz et al., 2020). For the experiment in Figure 1, dark-adapted (30 min) leaves were exposed to a light regime fluctuating between 1 min periods of 50 and 825 µmol photons m^-2^ s^-1^, with 250 ms dark intervals administered every 5 seconds for the first 30 seconds and every 10 seconds thereafter. For the methyl viologen (MV) experiment, detached leaves were dark-incubated overnight on water with 0.05% Tween-20 ± 1 µM MV. In the ECS measurement, dark intervals of 450 ms were administered every 4 s until 28 s during 95 µmol photons m^-2^s^-1^ of actinic illumination, every 10 s between 28 and 58 s, every 30 s between 58 and 298 s, every 4 s after transition to 825 µmol photons m^-2^s^-1^ between 298 and 318 s, and every 30 s thereafter. 2,000 Hz pulse frequency of the measuring light was used. ΔECS_T_ values were normalised to the ΔECS measured upon a 20 µs single-turnover saturating flash administered with the Dual-PAM 100 on dark-adapted leaves (ΔECS_ST_) (Mathiot & Alric, 2021). For the NEM experiment, detached leaves were incubated in dark on water with 0.05% Tween-20 ±0.1 mM NEM for 2 hours before the experiment. *Chlamydomonas* cultures and *S. viridis* leaves were measured using the same protocol as *Arabidopsis*, with *Chlamydomonas* chlorophyll concentration set to 20 Chl µg ml^-1^.

**Figure 1.**
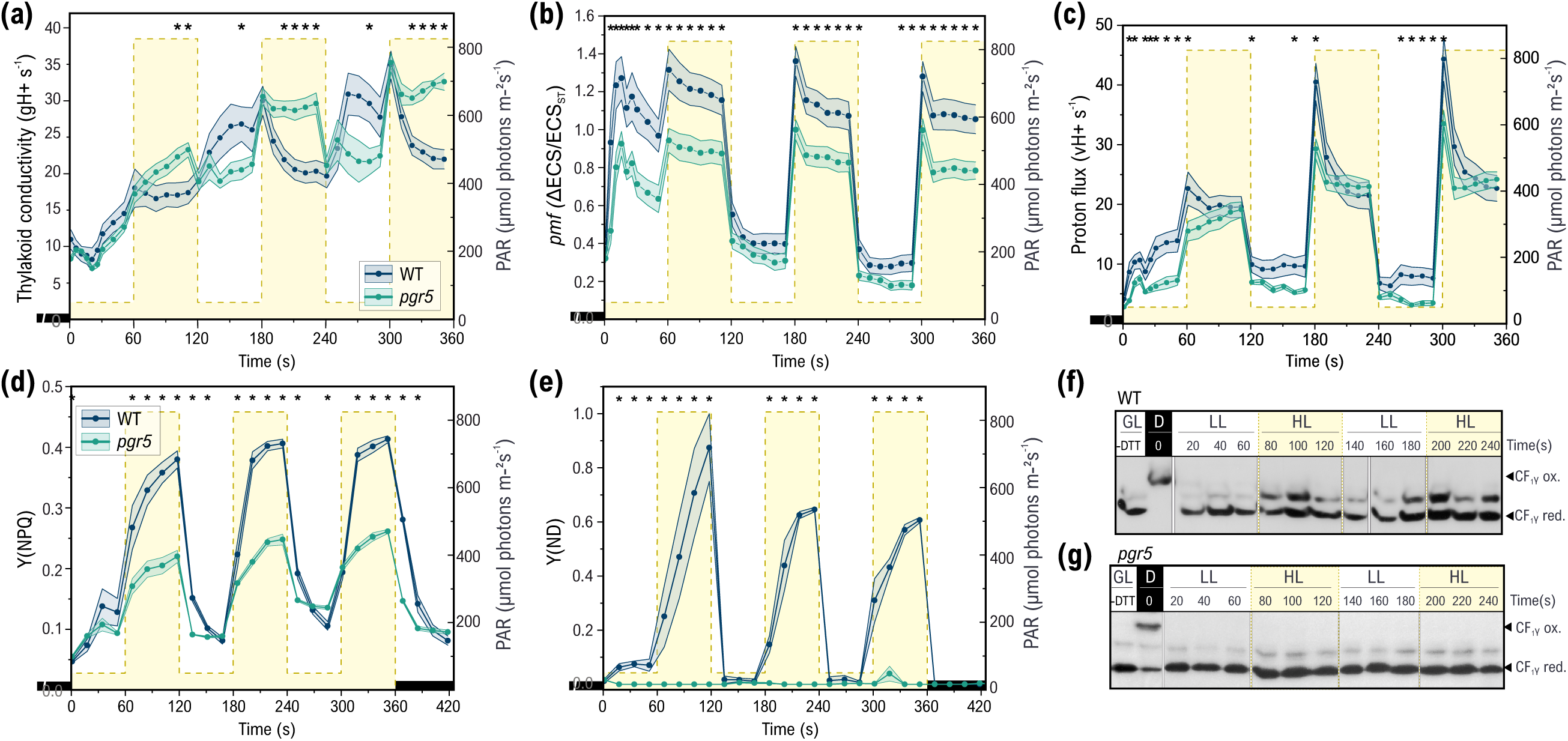
**Effect of PGR5 and CF_1_**_γ_ redox state on *pmf,* thylakoid conductivity, and photoprotection in fluctuating light in *Arabidopsis*. (a) Proton conductivity of the thylakoid membrane (g_H_+ s^-1^), **(b)** magnitude of the *pmf,* and **(c)** Thylakoid proton flux (v_H_+ = g_H_+ x *pmf*) in fluctuating light, as determined from dark-interval relaxation kinetics (DIRK) of the ECS signal. ECS was measured from dark-adapted WT-gl1 and *pgr5-1* mutant *Arabidopsis* leaves under a light regime alternating between 1 min periods of low (50 µmol photons m^-2^s^-1^) and high light (825 µmol photons m^-2^s^-1^). The values in (a) and (c) are averages from 9 and in (b) 11 biological replicates ± SEM. **(d)** Quantum yield of non-photochemical quenching (Y(NPQ)) and **(e)** fraction of oxidised P700 (donor side limitation of PSI, Y(ND)) in a fluctuating light regime supplemented with saturating pulses at 15 s intervals. The values in (d–e) are averages from 3 biological replicates ± SEM. In (a–e) * indicates statistically significant difference between WT and *pgr5-1* according to a two-sample Student’s t-test (P<0.05). **(f-g)** Mobility shift assays of the *in vivo* thiol redox state of the CF_1_y subunit of the chloroplast ATP synthase in WT (f) and in *pgr5-1* (g). Samples were taken from fluctuating light (1 min periods of 50 µmol photons m^-2^s^-1^, LL, and 825 µmol photons m^-2^s^-1^, HL. The break between the 140 and 160 sec samples in (f) represents division between two gels. –DTT indicates control sample where DTT was not added after NEM incubation and no alkylation of thiols by MALPEG should occur.

ECS changes were measured from *Synechocystis* and *Synechococcus* cells by monitoring the 500-480 nm absorbance difference signal (Viola et al., 2019) using a JTS-10 spectrophotometer (BioLogic) as described in (Nikkanen et al., 2020). Cells were pelleted and resuspended in fresh BG-11 at 7.5 µg Chl ml^-1^. When appropriate, 0.1 mM NEM was added to the samples before a 5 min dark-adaption prior to the measurement. Each measurement was also conducted with the cells illuminated only by the measuring light flashes, and the ECS changes recorded were then subtracted from the measurements under green actinic light, in order to control for any actinic effects of the measuring light. A moderate (60 µmol photons m^-2^ s^-1^, relative to growth light of 50 µmol photons m^-2^ s^-1^) rather than a low actinic light intensity was used in fluctuating light to allow a sufficient signal-to-noise ratio for dark-interval relaxation kinetics (DIRK) analysis. The cyanobacterial ECS signals were not normalised to ECS_ST_ due to a low signal-to-noise ratio of single turnover traces. Instead, a constant chlorophyll concentration was used.

The *pmf* and g_H_+ parameters were determined as the difference between magnitude of the ECS signal in light – y0 value of a first-order fit to the decay kinetics of the signal during a dark interval, and the inverse of the time constant of the fit, respectively, while v_H_+ was derived as *pmf* x g_H_+ (Cruz et al., 2005). Fitting and data processing was done with the OriginPro 2024b (OriginLab) software.

### Saturating pulse analysis of chlorophyll fluorescence and P700^+^

The extents of the donor side and acceptor side limitations of PSI, Y(ND) and Y(NA), respectively, were determined under fluctuating light conditions by a saturating pulse analysis of the P700^+^ signal using the Dual-PAM 100 (Klughammer & Schreiber, 2008). Chlorophyll autofluorescence was simultaneously recorded to determine the Y(NPQ) parameter (in *Arabidopsis*) as described by (Schreiber et al., 2008). Saturating pulses (8000 μmol photons m^−2^Js^−1^for *Arabidopsis* and 3000 μmol photons m^−2^Js^−1^ for *Synechocystis*) were administered every 15 seconds during illumination.

### Deconvoluted P700, plastocyanin, and ferredoxin redox changes

The deconvoluted plastocyanin (PC), P700, and ferredoxin redox changes were determined by NIR absorbance difference spectroscopy using the DUAL-KLAS-NIR spectrophotometer (Walz) (Klughammer & Schreiber, 2016). CET and LET at PSI in *Synechocystis* were quantified using DIRK analysis according to (Theune et al., 2021) with modifications. *Synechocystis* cells were pelleted and adjusted to 15 µg Chl ml^-1^ with fresh BG-11. Cells were pre-illuminated for 2 min with 500 μmol photons m^−2^□s^−1^, followed by administration of 100 dark intervals of 20 ms. The 100 decay traces were averaged and initial slopes of the PC and P700 decay curves were used to calculate J_PSI_. J_PSI_-CET was determined by repeating the same experiment in the presence of 10µM DCMU to block electron transport from PSII. The NIRMAX script (Klughammer & Schreiber, 2016) modified for cyanobacteria was used to determine the maximal oxidation of PC and P700 and maximal reduction of ferredoxin (Fed). The model spectra for the deconvolution of PC, P700, and Fed redox changes in *Synechocystis* were measured as described in (Nikkanen et al., 2020). Arabidopsis NIRMAX measurements of dark-adapted leaves were conducted as described in (Nikkanen et al., 2019). Reduction kinetics of Fed under far red light were measured after running the NIRMAX script, illuminating detached leaves for 1 second with intensity 20 far-red light.

### Cytochrome *f* redox changes

Detached leaves from 4-weeks-old WT-gl1 and *pgr5-1* Arabidopsis were dark-adapted for 5 min, after which the cytochrome *f* redox kinetics were measured as time-resolved absorbance changes at 554 nm with a baseline drawn between 546 and 573 nm during and after five seconds of illumination with 500 µmol photons m^-2^ s^-1^ of green light, using the JTS-10 spectrophotometer and sequential measurements with appropriate interference filters (10 nm FWHM, Edmund Optics). Scattering artifacts were minimised by BG39 filters (Schott). The cytochrome *f* post-illumination re-reduction rates (*k_f_* _red_ s^-1^) were obtained by fitting the post-illumination kinetics over 300 ms to a first-order exponential function.

### Protein thiol alkylation and SDS-PAGE

4-week-old *Arabidopsis* WT-Col-0 and *pgr5*-Cas plants were dark-adapted, exposed to a light regime fluctuating between 1-minute periods of 50 and 800 µmol photons m^−2^□s^−1^. Samples (individual leaves) were taken every 20 seconds and quickly placed in 10 % TCA solution to freeze the thiol redox state of proteins. TCA precipitation, alkylation of thiols with MALPEG-5000, and SDS-PAGE was then performed as described in Nikkanen et al., (2016). Briefly, free thiol groups were first alkylated with NEM. Following alkylation, disulphide bonds in proteins were reduced with dithiothreitol (DTT), and the resulting thiols were then alkylated using methoxypolyethylene glycol maleimide Mn 5000 (MAL-PEG, Sigma-Aldrich). This treatment makes the in-vivo oxidised forms of proteins migrate more slowly than reduced ones in SDS-PAGE. MV-treated WT-Col-0 leaves (see above) were illuminated in growth conditions for 0, 30, 90, or 180 min before sample collection and placement in 10% TCA.

Total protein extractions from *Synechocystis* cells were done as described in (Zhang et al., 2009) and from *N. benthamiana* leaves by grinding the tissue in a buffer with 100 mM Tris-HCl pH 8, 50 mM EDTA, 250 mM NaCl, and 0.75% SDS, heating for 10 min at 68°C, and centrifuging for 10 min at 14,000 rpm. Protein concentration was determined by the DC Protein Assay Kit (Bio-Rad, based on the Lowry assay). Sodium dodecyl sulphate polyacrylamide gel electrophoresis (SDS-PAGE) and immunoblotting were performed as previously described (Nikkanen et al., 2016). 4–15% Mini-PROTEAN TGX precast gels (Bio-Rad) were used for electrophoresis except for Figure 5, for which 12% polyacrylamide gels with 6M urea were cast. PVDF membranes were probed with antibodies raised against CF_1_γ (Agrisera, AS08 312), FBPase (courtesy of Dr M. Sahrawy), AtpB (Agrisera, AS), Flv3 (Genscript), NTRC (Lepistö et al., 2009), or 2-CysPrx (courtesy of Prof. F.JJ. Cejudo).

### Membrane Inlet Mass Spectrometry (MIMS)

*Synechocystis in vivo* O_2_ and CO_2_ gas fluxes were monitored in real-time by membrane inlet mass spectrometry (MIMS) as described earlier (Mustila et al., 2016), with O_2_ uptake distinguished from O_2_ evolution by enrichment of the ^18^O_2_ isotope in the medium. Chlorophyll concentration was set to 10 µg Chl ml^-1^, 1.5 mM NaHCO_3_ was added, and cells were dark-adapted for 15 min prior to the measurements. Gas exchange was monitored under 500 μmol photons m^-2^s^-1^ illumination for 5 min. See legend for Fig. 2i for experimental details.

**Figure 2.**
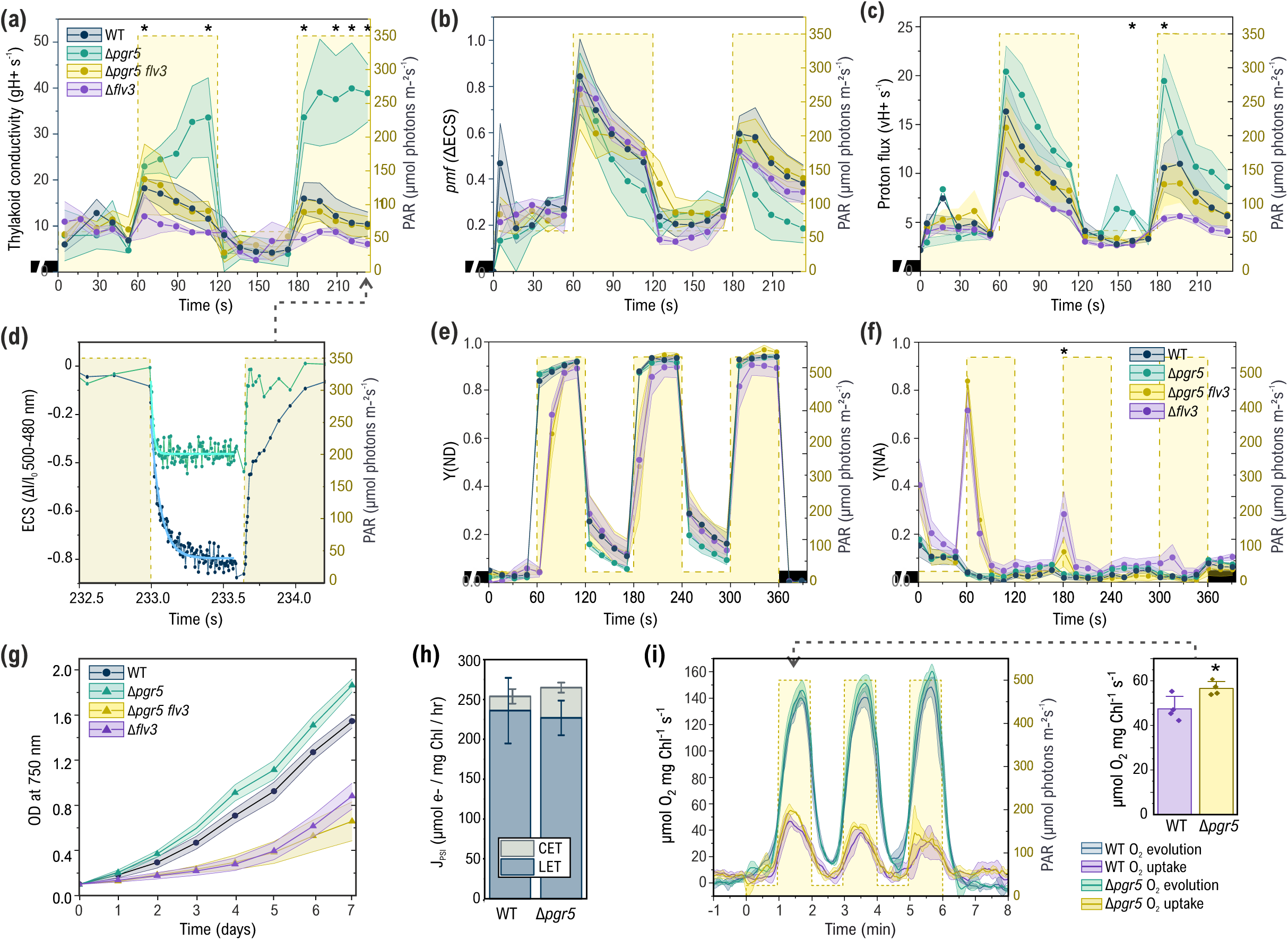
***Pmf* dynamics, PSI oxidation, O_2_ fluxes, and growth under fluctuating light in *Synechocystis*.** Thylakoid conductivity (g_H_+) **(a)**, *pmf* **(b),** and proton flux (vH+) **(c)** were determined by the ECS DIRK method from WT, Δ*pgr5,* Δ*pgr5 flv3,* and Δ*flv3* cells. The values in a–c are averages from 3–6 (depending on the quality of the first-order DIRK fit) biological replicates ± SEM, apart from the second data points of WT and Δ*pgr5*, where signal-to-noise ratio allowed only single replicates to be fitted successfully. * indicates statistically significant differences between WT and Δ*pgr5* according to a two-sample Student’s t-test (P<0.05). **(d)** Representative ECS decay kinetics in WT and Δ*pgr5* cells during the second-to-last 600 ms dark interval in the experiment. First order exponential fits are shown in cyan and turquoise. **(e)** Donor (Y(ND)) and **(f)** acceptor side limitation (Y(NA)) of PSI in WT, Δ*pgr5,* Δ*pgr5 flv3*, and Δ*flv3* strains of *Synechocystis* during transitions from low light (25 µmol photons m^-2^s^-1^, LL) to high light (525 µmol photons m^-2^s^-1^, HL). Y(NA) and Y(ND) were determined from P700 redox kinetics using the saturating pulse method on a DUAL-PAM 100 spectrophotometer. Traces are averages of 3–4 biological replicates ± SEM (ribbons). * indicates statistically significant difference to WT according to a two-sample Student’s t-test (P<0.05). **(g)** Growth curves (OD 750 nm) of WT, Δ*pgr5* and Δ*pgr5 flv3* cells under fluctuating light (5 min at 20 µmol photons m^-2^s^-1^ to 30 sec at 500 µmol photons m^-2^s^-1^). The values are averages from 3 biological replicates ± SEM. **(h)** Quantification of LET and CET rates in *Synechocystis* WT and Δ*pgr5*. Chlorophyll was adjusted to 20 µg Chl ml^-1^ in BG-11 pH 7.5. Using a Dual KLAS-NIR spectrophotometer, cells were illuminated for 2 min under 500 µmol photons m^-2^s^-1^before administering 100x 40 ms dark intervals 10 s apart to monitor the relaxation kinetics of the P700 and PC signals. The measurement was then repeated in the presence of 10µM DCMU to inhibit LET. Values means of four biological replicates ± SD. **(i)** O_2_ gas fluxes in fluctuating light in WT and Δ*pgr5* as measured by membrane inlet mass spectrometry (MIMS). Cells were grown in fluctuating light (1 min 25 µmol photons m^-2^s^-1^ / 1 min 500 µmol photons m^-2^s^-1^), adjusted to 10 µg Chl ml^-1^, and ^18^O_2_ was added in equilibrium with ^16^O_2_ in order to distinguish O_2_ uptake from O_2_ evolution. Cells were dark-adapted for 10 min, after which real-time gas fluxes were measured by MIMS mimicking the fluctuating growth light conditions, with a 5 min dark period on either side of three LL-HL cycles. The values are averages of four biological replicates ± SD. The bar chart shows the maximum O_2_ uptake rates + SD during the first HL phase of the experiment (dotted arrow). * Indicates statistically significant difference between WT and Δ*pgr5* according to a two-sample Student’s t-test (P<0.05).

### Bimolecular Fluorescence Complementation (BiFC) tests

*In planta* protein–protein interaction tests by bimolecular fluorescence complementation (BiFC) were conducted as described earlier (Nikkanen et al., 2016, 2018). *Nicotiana benthamiana* plants were used for transient expression of fusion constructs via *Agrobacterium tumefaciens* strain GV3101 infiltration. Imaging was done 3 days after infiltration using a Zeiss LSM780 confocal microscope.

Construction of the AtPGR5:YFP-C and AtNdhS:YFP-N expression vectors for YFP-fusion proteins is described in (Nikkanen et al., 2018) and construction of the AtCF_1_γ:YFP-N and AtTRXx:YFP-C vectors in (Nikkanen et al., 2016).

### In silico modelling

The ClusPro 2.0 protein-protein docking service (Kozakov et al., 2017) was used to perform molecular docking of the AtPGR5 (Uniprot identifier Q9SL05) / SoCF1γ (P05435) / SoCF1□ (P00833) interaction complex, as well as the AtPGR5 / AtPGRL1a (Q8H112)/ AtTRXm4 (Q9SEU6) complex. The predicted chloroplast transit peptides (amino acid residues 1–60, 1–60, and 1–82) for AtPGR5, AtPGRL1a, and AtTRXm4, respectively) were removed from the structures before docking.

The amino acid sequences for *Synechocystis* genes *sll1327, slr1330* and *ssr2016* were retrieved from the Uniprot database (entries P17253, P26533, and P73358, respectively). AlphaFold2 Multimer Google Colab notebook (Mirdita et al., 2022) was used to model the interaction complex using 6 recycles and pdb100 template mode, with Amber relaxation of the best model. The model with the best pTM score was selected for visualisation using the Mol* 3D viewer (Sehnal et al., 2021). The *Spinacia oleracea* chloroplast ATP synthase structure (Hahn et al., 2018) was retrieved from the

RCSB PDB database (entry 6FKF), while overlaying with the SoCF1γ/AtPGR5 interaction model as well as modelling of electrostatic surface charges was performed with ChimeraX (Meng et al., 2023).

### Statistical tests

ANOVA, Tukey’s tests for significance, Student’s T-tests, and linear mixed effects modelling (LMEM) were performed using the OriginPro 2024 software and its LMEM App. The LMEM models included gH□ as the dependent variable, *pmf* as a continuous covariate, genotype and light intensity as fixed factors, and biological replicate as a random factor. Significance was assessed using the F-tests provided by the software.

## RESULTS

### Dynamic changes in thylakoid conductivity are independent of the thiol redox state of CF_1_**_γ_** in plants

In plants and algae, measurement of dark-interval relaxation kinetics (DIRK) of the electrochromic shift (ECS) signal has been widely used for *in vivo* determination of the magnitude of the *pmf*, partitioning of the *pmf* between ΔpH and ΔΨ, as well as the conductivity of the thylakoid membrane, which is mainly determined by the activity of ATP synthase (Sacksteder & Kramer, 2000; Cruz et al., 2001, 2005; Schreiber & Klughammer, 2008; Bailleul et al., 2010). Recently, a reliable method to measure ECS has also been developed for cyanobacteria (Viola et al., 2019; Nikkanen et al., 2020; Santana-Sánchez et al., 2023).

Time-resolved ECS analysis in wild type (WT) *Arabidopsis* revealed that upon the onset of illumination of dark-adapted leaves, the conductivity of the ATP synthase (g_H_+) initially decreased for c.a. 30 s (Fig. 1a), as observed previously (Nikkanen et al., 2018; Guinea Diaz et al., 2020). This initial decrease in g_H_+ allowed transient build-up of *pmf* (Fig. 1b) and induction of NPQ and photosynthetic control (as indicated by donor-side limitation of PSI) (Fig. 1d–e) in order to avoid over-reduction of the electron transport chain when redox-regulated CBC enzymes are not yet activated (i.e. reduced).

After the initial decrease, at c.a. 30 s of illumination, g_H_+ started to increase gradually, resulting in decrease of *pmf* (Fig. 1a–d). Importantly, these changes in g_H_+ did not correlate with the thiol redox state of CF_1_γ. CF_1_γ was rapidly reduced at the onset of illumination, becoming almost fully reduced after 20 s, and remained reduced thereafter (Fig. 1f). In contrast, the CBC enzyme Fructose-1,6-bisphosphatase (FBPase) remained mostly oxidised (inactive) for a full minute in low light, and only became partially reduced when leaves were exposed to HL (Supplementary Fig. S2).

In a light regime where light intensity alternates between periods of low light (LL) and HL, mimicking natural fluctuations, intricate regulatory dynamics of the *pmf* and g_H_+ could be observed. During LL phases, g_H_+ increased substantially, which contributed to a rapid decline in *pmf* (Fig. 1a– b). This upregulation of g_H_+ allowed relaxation of photosynthetic control (as inferred from donor-side limitation of PSI) and NPQ (Fig. 1e–f), allowing sufficient supply of electrons to stromal reactions.

Similarly to downregulation of g_H_+ during dark-to-light transitions, the upregulation of g_H_+ during LL happened independently of the thiol redox state of CF_1_γ, which remained fully reduced throughout the change in light intensity (Fig. 1e).

In contrast to LL phases, g_H_+ was rapidly downregulated in WT leaves during HL phases (Fig. 1a), enabling sufficient *pmf* build-up (Fig. 1b) for induction of photosynthetic control and NPQ (Fig. 1d– e). Again, the decrease in g_H_+ did not coincide with oxidation of CF_1_γ, which remained fully reduced (Fig. 1f).

These results demonstrate that ATP synthase activity undergoes dynamic regulation under fluctuating light in Arabidopsis, and that this regulation is independent of CF_1_γ redox state and must occur via another mechanism.

### Thylakoid conductivity is dynamically regulated under fluctuating light also in cyanobacteria and unicellular algae

When WT *Synechocystis* cells were subjected to sudden fluctuations between moderate light (ML) and HL, time-resolved ECS-DIRK measurements revealed downregulation of g_H_+ during HL phases (Fig. 2a), resembling that observed in *Arabidopsis*. As in plants, this downregulation likely occurs to maintain an adequate *pmf* (Fig. 2b) for induction of photosynthetic control. Accordingly, strong donor side limitation of PSI (Y(ND)) was measured during HL phases (Fig. 2e) indicative of limitation of electron transport from Cyt b_6_f. In contrast to *Arabidopsis*, g_H_+ remained low during the ML phases of the experiment (Fig. 2a).

We also probed the *pmf* and g_H_+ dynamics under fluctuating light in the unicellular green alga *Chlamydomonas*. While *pmf* underwent similar changes as in plants and in cyanobacteria (Fig. 3b), no clear downregulation of g_H_+ was observed during HL, and during the LL phases g_H_+ remained low (Fig. 3a).

**Figure 3.**
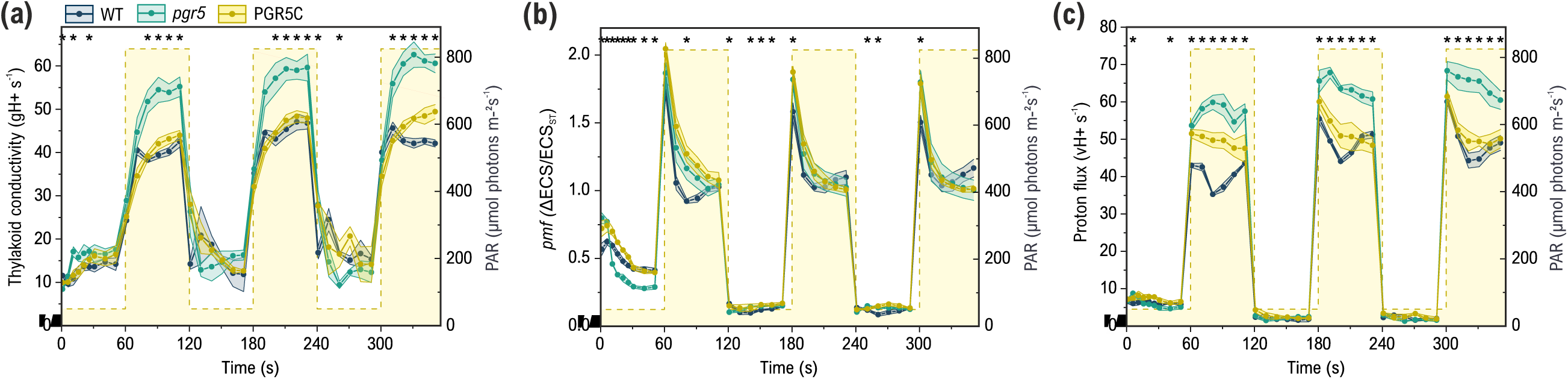
*Pmf* dynamics in *Chlamydomonas reinhardtii* under fluctuating light. **(a)** Thylakoid conductivity (g_H_+). **(b)** *pmf* (ΔECS/ECS_ST_). **(c)** Proton flux (v_H_+). The ECS signal was measured from WT137c, *pgr5* knockout mutant, and PGR5 complementation line (PGR5C) cells under a fluctuating light regime, alternating between 1 min periods of low light (50 µmol photons m^-2^s^-1^) and high light 825 (µmol photons m^-2^s^-1^). Values are means ± SEM from three biological replicates. The asterisks indicate statistically significant differences between WT and *pgr5* according to a two-sample Student’s t-tests (P<0.05).

### PGR5 is needed for downregulation of thylakoid conductivity under strong illumination across photosynthetic organisms

During dark-to-LL transitions, the *Arabidopsis pgr5* mutant plants had a lower *pmf* than WT (Fig. 1B), as shown previously and attributed to lower linear electron transport rates (Nikkanen et al., 2018). In fluctuating light, the *pgr5* mutants maintained a significantly higher gH+ than WT during HL phases (Fig. 1a), resulting in partial loss of *pmf* (Fig. 1b), in agreement with previous studies (Avenson et al., 2005; Nikkanen et al., 2018). The mutant plants also exhibited impaired induction of NPQ (Fig. 1d) and photosynthetic control, resulting in an inability to oxidise P700 (Y(ND), Fig. 1e) and low PSI quantum yield (Supplementary Fig. S3a) due to high acceptor side limitation (Supplementary Fig. S3b). Interestingly, and in contrast to their behaviour during HL phases, *pgr5* mutant plants showed a lower g_H_+ and slightly lower *pmf* than WT during LL phases (Fig. 1a–b), which was due to diminished proton flux (vH+) (Fig. 1c). A small amount of reduced CF_1_γ was detected in the dark in *pgr5*, but under the fluctuating light treatment the redox state of CF_1_γ was indistinguishable from WT, as CF_1_γ was rapidly reduced upon illumination and maintained as such throughout the fluctuating light treatment also in *pgr5* (Fig. 1g).

As it has been proposed that PGR5 deficiency results in impaired Q cycle activity in *Chlamydomonas* (Buchert et al., 2020), we confirmed that the elevated gH+ values in Arabidopsis *pgr5* are not caused by diminished charge separation in the Q cycle during the dark intervals. No difference in post-illumination re-reduction rates of cytochrome *f* was detected between WT and *pgr5* (Supplementary Fig. S4), indicating that the elevated gH+ in *pgr5* does indeed derive from increased proton conductivity of the thylakoid membrane.

Next, we sought to elucidate the bottleneck in the electron transport chain that impairs PSI oxidation in *pgr5*. It was proposed recently that Fed binding to PSI is impaired due to damage of F_A_ and F_B_ iron-sulphur clusters in *pgr5* already under moderate light (Tiwari et al., 2024). Since it has also been suggested that the rate of Fed oxidation is primarily determined by the linear electron transport rate (Takagi & Miyake, 2018), we compared the Fed reduction and oxidation kinetics between WT and *pgr5* during and after a 1 s pulse of far-red light to minimise the contribution of linear electron flow and examine just the ability of PSI to reduce, and acceptors to oxidise Fed. Our results show that both reduction and re-oxidation kinetics of Fed are slowed down in *pgr5,* while oxidation of PC was increased and oxidation of P700 remained similar to WT (Supplementary Fig. S5), in agreement with the proposal by Tiwari et al. (2024).

The *Arabidopsis pgr5-1* mutant used in our experiments has been reported to also contain an additional mutation in the At2g1724 gene, which slightly exacerbates PSI photoinhibition in the mutant (Wada et al., 2021). Mutants where an intact At2g1724 has been re-introduced (*pgr5*^hope1^ and *pgr5-Cas*) are, however, also unable to induce photosynthetic control to oxidise PSI (Wada et al., 2021), and have similarly elevated g_H_+ and diminished *pmf* as the *pgr5-1* mutant (Penzler et al., 2022; Maekawa et al., 2024). While it is therefore unlikely that the At2g1724 mutation in *pgr5-1* plants would have a significant effect on the phenotypes examined in the current study, we confirmed this by repeating the ECS experiment (Fig. 1a–c) also with the *pgr5-Cas* mutant (Penzler et al., 2022).

Results with *pgr5-Cas* plants were similar to the ones with *pgr5-1,* apart from the decrease in *pmf* during initial dark-to-LL transitions being less prominent in *pgr5-Cas* plants than in *pgr5-1* in comparison to WT (Supplementary Fig. S6). Moreover, while a diminished proton flux (v_H_+) in LL and immediately upon shifts to HL was measured in *pgr5-1* mutants in comparison to WT (Fig. 1c), this decrease was absent in *pgr5-Cas* plants, which instead exhibited elevated v_H_+ during the first HL phase (Supplementary Fig. S6c). In order to examine if this elevation of vH+ or changes to gH+ were linked to altered mitochondrial respiration, we repeated the ECS experiment in the presence of potassium cyanide (KCN) and salicylhydroxamic acid (SHAM), inhibitors of Complex IV and alternative oxidase (AOX) in mitochondrial respiration, respectively. Compared to WT, *pgr5-*Cas plants maintained elevated gH+ also in these conditions, while vH+ and *pmf* are lower than WT in LL but higher than WT in HL (Supplementary Fig. S7). These results indicate that the elevated gH+ and vH+ observed in HL in *pgr5* mutants do not derive from increased mitochondrial respiration.

We proceeded to investigate whether lack of SynPgr5 affects regulation of g_H_+ in *Synechocystis*. Elevated g_H_+ was indeed detected upon exposure of Δ*pgr5* cells to HL after illumination in moderate light (ML) in a fluctuating light regime (Fig. 2a, d), concomitant with lowered (albeit statistically insignificantly) *pmf* towards the end of the second HL cycle (Fig. 2b). The increased g_H_+ was not due to an elevated amount of ATP synthase, as the protein level of AtpB remained unchanged in the mutant (Supplementary Fig. S8). Despite the elevated g_H_+ and in contrast to AtPGR5-deficient

*Arabidopsis*, no decrease in PSI quantum yield (Supplementary Fig. S3c), donor-side limitation (indicative of impairment of photosynthetic control) (Fig. 2e) or increase in acceptor side limitation of PSI (Fig. 2f) was detected during transitions to HL in *Synechocystis* Δ*pgr5* cells. Interestingly, a slight, albeit statistically non-significant decrease in donor side limitation of PSI was detected in Δ*pgr5* upon transitions from HL to LL (Fig. 2e).

The ability of the *Synechocystis* Δ*pgr5* strain to maintain sufficient PSI oxidation for unimpaired photosynthetic activity may be due to the strong electron sink capacity of the Mehler-like reaction catalysed by flavodiiron proteins (FDPs) under light (Allahverdiyeva et al., 2013). Therefore, we tested the PSI oxidation capacity of a Δ*flv3 pgr5* double mutant strain, which is deficient in both SynPgr5 as well as the Flv1/3 hetero-oligomers (Supplementary Fig. S1) which are mainly responsible for the strong, rapidly induced Mehler-like reaction (Santana-Sanchez et al., 2019). We detected slightly but non-significantly aggravated acceptor side limitation of in the double mutant in comparison to the Δ*flv3* single mutant during initial LL-to-HL transitions (Fig. 2f). The additional effect of Pgr5 deficiency was minor compared to lack of Flv3, and no additional aggravation of ferredoxin redox kinetics, CO_2_ fixation, or O_2_ evolution was observed in Δ*flv3 pgr5* in comparison to Δ*flv3* (Supplementary Fig. S9). These results indicate that SynPgr5 may affect PSI oxidation also in cyanobacteria, but its importance is minor compared to plants, while flavodiiron proteins have a crucial role. This was also reflected in the growth phenotypes of the strains. In agreement with previous results (Allahverdiyeva et al., 2013), no impairment of growth was detected between WT and Δ*pgr5* under fluctuating light (1 min LL / 1 min HL), with Δ*pgr5* even growing slightly faster than WT, while additional deficiency of Flv3 in Δ*pgr5 flv3* and Δ*flv3* resulted in severe impairment of growth (Fig. 2g).

The Δ*flv3* mutant showed a decreased g_H_+ in the HL phases of fluctuating light (Fig. 2a), which allowed it to compensate for diminished v_H_+ (Fig. 2c) and maintain WT-level *pmf* (Fig. 2b). The Δ*pgr5 flv3* double mutant, however, exhibited WT-like g_H_+, v_H_+, and *pmf* (Fig. 2a–c), indicating that the downregulation of g_H_+ observed in Δ*flv3* is also at least partly dependent on SynPgr5.

To clarify whether SynPgr5 is involved in CET in *Synechocystis* as previously concluded (Yeremenko et al., 2005; Dann & Leister, 2019), we quantified the rates of CET and linear electron transport (LET) in WT and Δ*pgr5* cells using DIRK analysis of near-infrared absorbance difference signals, as described by (Theune et al., 2021). Our results showed that the CET rate was not decreased in the deletion mutant (Fig. 2h), indicating that SynPgr5 is not involved in CET, at least in steady state conditions. It has been previously reported that a *Synechocystis* Δ*pgr5* mutant has an accelerated rate of P700 oxidation under far-red light as measured with the Dual-PAM 100 spectrophotometer (Dann & Leister, 2019), which can be an indication of decreased CET, although results from such assays should be interpreted with caution (Fan et al., 2007). In contrast to the previous report, we detected no difference between WT and Δ*pgr5* in far-red-induced P700 oxidation rate when measured as the deconvoluted P700 signal using the Dual-KLAS-NIR instrument (Supplementary Fig. S10). There results suggest that SynPgr5 is not involved in CET.

Near-infrared spectroscopy revealed no significant differences in the deconvoluted redox signals for ferredoxin (Fed), plastocyanin (PC) or P700 between WT and Δ*pgr5* cells upon onset of strong illumination (Supplementary Fig. S11a). Moreover, analysis of *in vivo* O_2_ and CO_2_ gas exchange by membrane inlet mass spectrometry (MIMS) showed that oxygen evolution (Supplementary Fig. S11b), dark respiration rate (Supplementary Fig. S11c), light-induced O_2_ uptake rate indicative of FDP activity (Supplementary Fig. S11b, d), as well as CO_2_ fixation (Supplementary Fig. S11e) were all unchanged in Δ*pgr5* in comparison to WT cells. When cells were grown under fluctuating light and O_2_ gas exchange was measured under similar conditions, however, a slightly elevated light-induced O_2_ uptake rate was observed in Δ*pgr5,* although the difference was statistically significant only during the first HL phase (Fig. 2i).

There is no PGR5 orthologue identified in the genome of another cyanobacterial model species, *Synechococcus elongatus* PCC 7942 (*Synechococcus*) (Shimakawa et al., 2016). We therefore proceeded to investigate whether the regulation of g_H_+ in response to fluctuating light conditions in *Synechococcus* differs from *Synechocystis*. Interestingly, no major difference was observed in comparison to the g_H_+ kinetics in *Synechocystis*, with down-regulation of g_H_+ occurring during the HL phases after an initial increase (Supplementary Fig. S12). This suggests that the downregulation of g_H_+ in HL is achieved via a different mechanism in *Synechococcus*.

Elevated g_H_+ under HL was also observed in the *Chlamydomonas pgr5* mutant, in comparison to WT or a CrPGR5 complementation strain (Fig. 3a). Interestingly, this did not result in lower *pmf* under HL due to elevated proton flux (v_H_+) (Fig. 3b, c). We confirmed that the potential presence of damaged PSI centres in *pgr5* (Tiwari et al., 2024) did not result in major overestimation of the *pmf* and vH+ parameters by also plotting the *Chlamydomonas* ECS data normalised only by Chl concentration (Supplementary Fig. S13). Finally, the increase in g_H_+ during the HL phases of a fluctuating light regime was also detected in a *pgr5* mutant of the model C_4_ grass *S. viridis* (Supplementary Fig. S12). Similarly to Arabidopsis but unlike *Synechocystis*, both *Chlamydomonas* (Jokel et al., 2018) and *S*. *viridis* (Woodford et al., 2024) *pgr5* mutants suffer from an inability to oxidise PSI in HL.

It has been suggested that the elevated gH+ in *pgr5* mutants is simply a consequence of lower *pmf*, and that when *pmf* is the same, there would be no difference in gH+ between WT and *pgr5* (Kappel et al., 2023). However, this correlation was made based on steady state measurements. When we plotted the gH+ values from our time-resolved ECS measurements in fluctuating light as a function of *pmf*, the correlation between the two parameters differed dramatically between WT and *pgr5* mutants in all species examined in the current study (Fig. 4). Linear mixed effects models (Supplementary Tables S1–S7) revealed that Arabidopsis *pgr5-1* mutants exhibited a significantly elevated gH□ in HL compared to WT independently of *pmf* (p=0.001), with an estimated difference of +5.6 gH+ units across all *pmf* values. Notably, *pmf* was not a significant predictor of gH□ in this condition (p=0.74). Similarly, gH+ values increased independently of *pmf* in HL also in Arabidopsis *pgr5*-Cas (+6.89, p=0.001), and in the full dataset in *Synechocystis* Δ*pgr5* (+12.3, p=0.0154), *Chlamydomonas pgr5* (+5.72, p=0.003, and *S. Viridis pgr5* (+10.7, p=0.010). These results indicate that the effect of PGR5 on gH+ is not an indirect consequence of diminished *pmf* in any of the studied organisms, and highlight a conserved function of PGR5 in regulation of the proton conductivity of ATP synthase from cyanobacteria to chloroplasts in algae, C_3_, and C_4_ plants.

**Figure 4.**
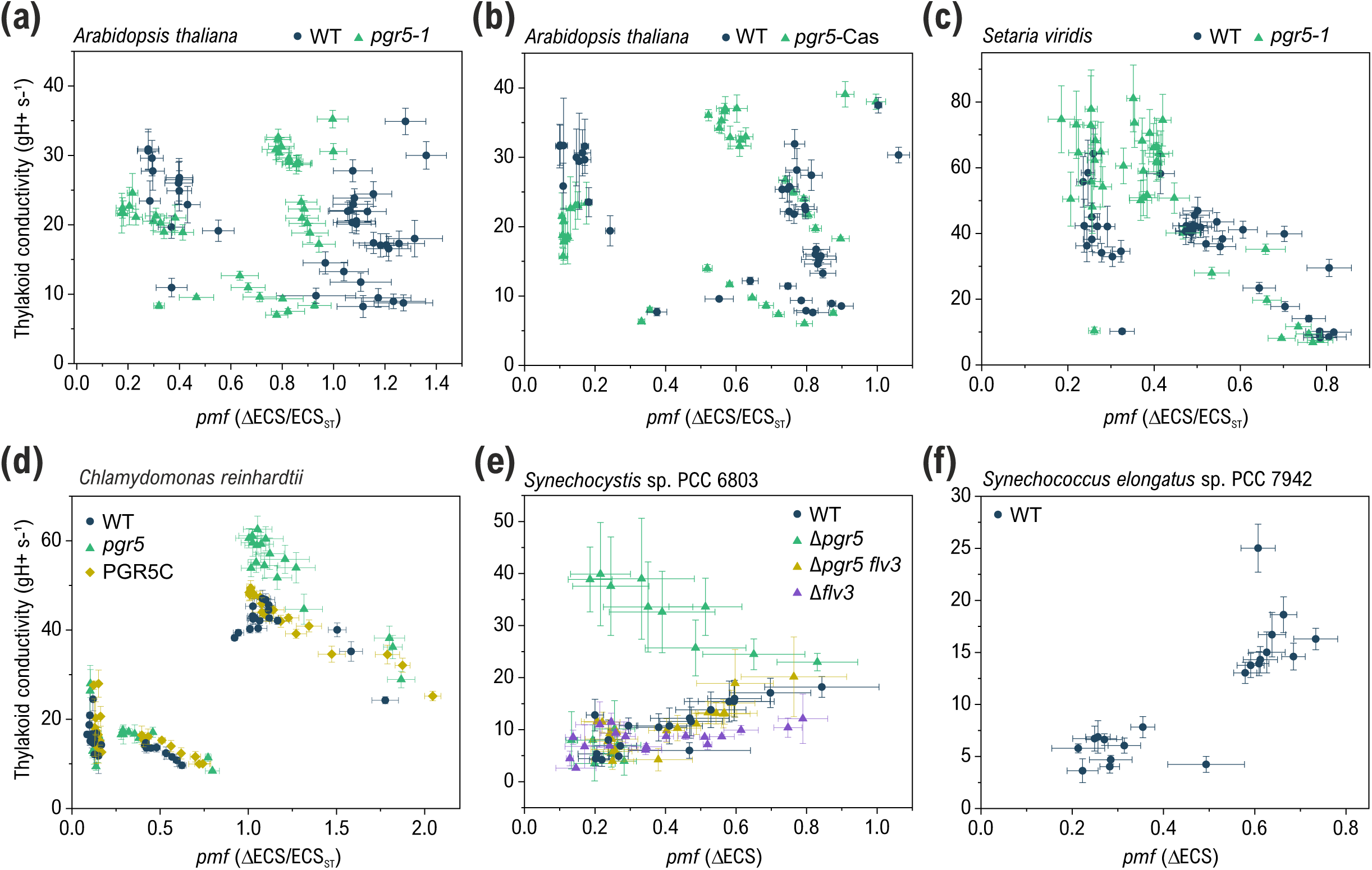
Correlation of *pmf* with gH+ in WT and PGR5-deficient mutants across photosynthetic organisms. Time-resolved ECS datasets from fluctuating light conditions. **(a)** *Arabidopsis thaliana* dataset from Fig. 1a–c. **(b)** *Arabidopsis thaliana* dataset from Supplementary Fig. S4. **(c)** *Setaria viridis* dataset from Supplementary Fig. S12. **(d)** *Chlamydomonas reinhardtii* dataset from Fig. 3. **(e)** *Synechocystis* sp. PCC 6804 dataset from Fig. 2a–c. (f) *Synechococcus elongatus* sp. PCC 7942 dataset from Supplementary Fig. S11.

**Figure 5.**
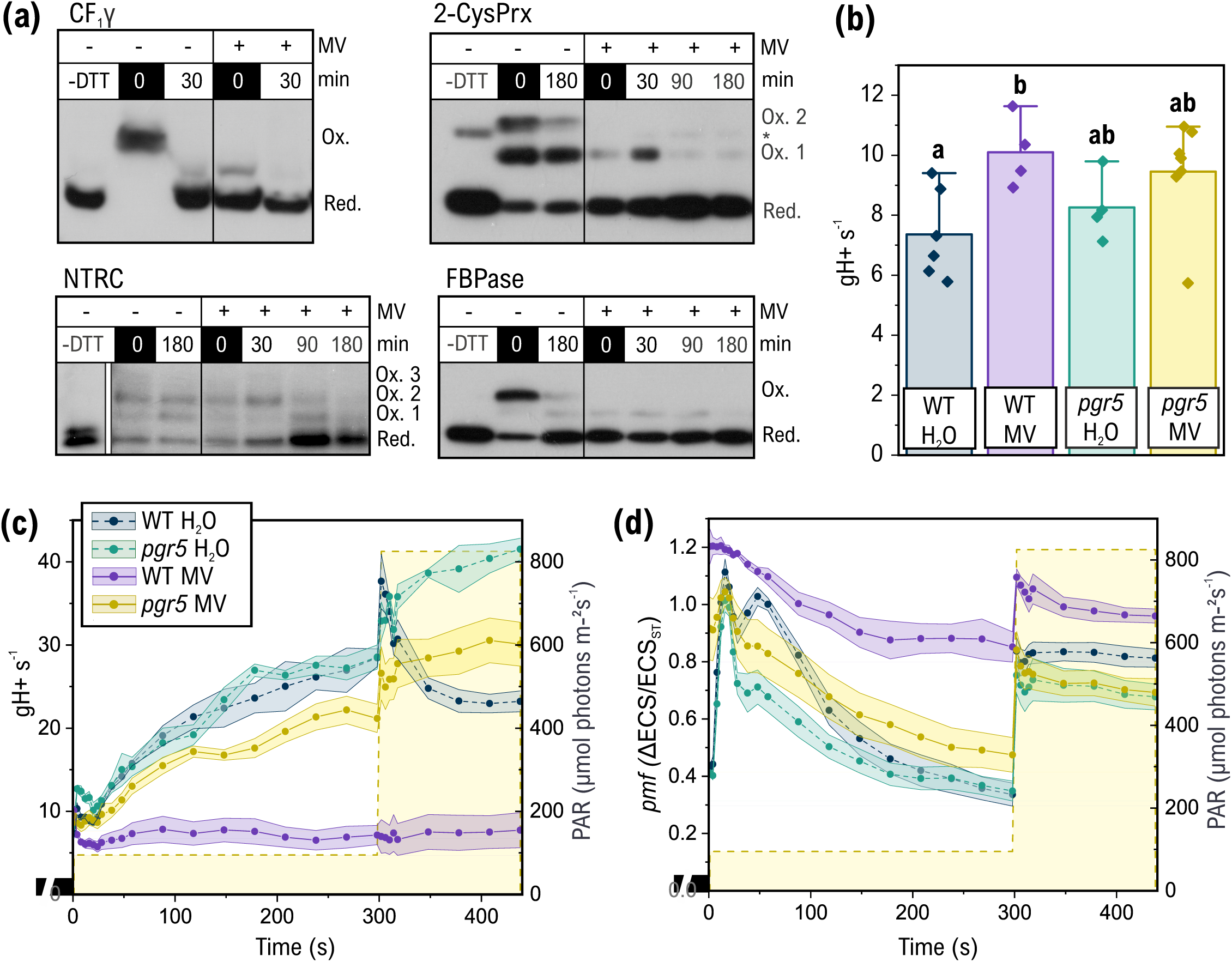
Effect of methyl viologen (MV) on thiol redox states of chloroplast enzymes and on regulation of *pmf* in *Arabidopsis*. **(a)** Mobility shift assays of AtCF_1_y, 2-CysPrxs, NTRC, and FBPase thiol redox state in vivo. WT Arabidopsis leaves were incubated in H_2_O with (+) or without (-) 1 µM MV in dark and then illuminated in GL conditions for 180 min, with samples taken from darkness (0) and after 30, 90, and 180 min of illumination. Total protein extracts were labelled with MALPEG as described in Materials and Methods, separated by SDS-PAGE and probed with appropriate antibodies. MALPEG-labelled (in vivo oxidised) forms migrate more slowly than unlabelled forms (in vivo reduced). –DTT indicates control sample where no alkylation of thiols by MALPEG should occur, and * indicates dimeric 2CysPrx. We cannot fully exclude the possibility of the apparent reduction of chloroplast enzymes in presence of MV being caused by overoxidation of thiols to sulphenic acids (Huang et al., 2019), especially in the case of 2-CysPrxs (Puerto-Galan et al., 2013; Cerveau et al., 2016). S-sulphenylated proteins cannot form disulphides and thus cannot be labelled by the alkylating agent in our protocol, migrating similarly to fully reduced proteins in mobility shift assays. Apart from 2-CysPrxs, where the thiols are required for the catalytic activity of the enzyme, overoxidation may in fact result in in a conformation that is functionally similar to the reduced state of the protein. **(b)** Conductivity of the thylakoid membrane (g_H_+) after 1 s of illumination in growth light (GL, 95 μmol photons m^-2^s^-1^) in dark-adapted WT and *pgr5* mutant leaves incubated in H_2_O with or without 1 µM MV. The values are mean + SD from 4–7 biological replicates. Statistical significance of differences was tested by one-way ANOVA and Tukey’s tests (p<0.05). **(c)** Thylakoid conductivity and **(d)** *pmf* in dark-adapted WT and *pgr5* mutant leaves incubated in H_2_O with or without 1 µM MV during 5 min illumination in GL (95 μmol photons m^-2^s^-1^) and consequent increase in light intensity to 825 μmol photons m^-2^s^-1^ (HL), as determined from dark-interval relaxation kinetics of the ECS signal. The values in C-D are averages from 4–7 biological replicates ± SEM (ribbon).

### Elucidating the mechanism of redox-dependent inhibition of thylakoid conductivity

We have previously shown that increasing the reducing capacity of the chloroplast thiol redox regulation systems through overexpression of NTRC in *Arabidopsis* intensifies the downregulation of g_H_+ upon increases in light intensity (Nikkanen et al., 2018). We therefore proceeded to investigate potential thiol regulation of thylakoid conductivity or ATP synthase activity, as well as the role of PGR5 in it.

Addition of methyl viologen (MV) results in lowered conductivity of the thylakoid membrane, which has been interpreted as prevention of TRX-mediated reduction of the CF_1_γ disulphide (Kramer & Crofts, 1989). MV facilitates electron transfer from PSI or reduced ferredoxin to O_2_ (the Mehler reaction), resulting in generation of ROS (Fujii et al., 1990; Sétif, 2015; Shapiguzov et al., 2019, 2020). However, MV is also able to donate electrons to redox-regulated enzymes such as nitrite reductase in spinach (Hirasawa & Tamura, 1980) and glutathione reductase in yeast (Llobell et al., 1986). To further elucidate the relationship between stromal/cytosolic thiol redox state and PGR5-dependent downregulation of thylakoid conductivity, we investigated the effects of MV on chloroplastic thiol-regulated enzymes and *pmf* in *Arabidopsis.* We first determined the *in vivo* thiol redox states of the chloroplast redox enzymes as well as AtCF_1_γ in presence or absence of 1 µM MV. Although electron supply to the TRX systems from PSI is limited by MV, addition of MV caused strong reduction of the 2-cysteine peroxiredoxin (2-CysPrx), FBPase, and CF_1_γ pools even in darkness (Fig. 5a). NTRC, whose redox state is mostly unaffected by light conditions, thereby acting as a buffer system for plastidial redox state (Nikkanen et al., 2018; Zimmer et al., 2021), also became strongly reduced after 90 min of illumination when treated with MV (Fig. 5a).

Next, we investigated the effect of MV on regulation of *pmf* and g_H_+ in WT *Arabidopsis* leaves. Upon illumination of dark-adapted leaves, g_H_+ was initially significantly higher in MV-treated WT leaves than in control leaves (Fig. 5b), most likely due to dark-reduction of AtCF_1_γ by MV (Fig. 5a). As illumination continued, g_H_+ diminished in MV-treated leaves and remained at a very low level throughout the rest of the experiment (Fig. 5c). This results in very high *pmf* (Fig. 5d) and NPQ (Shapiguzov et al., 2020). In contrast, MV-treatment of *pgr5* mutant leaves caused only a minor decrease in g_H_+ and increase in *pmf* during illumination, and in HL g_H_+ was in fact higher in MV-treated *pgr5* mutant than in WT control leaves (Fig. 5c–d), suggesting that the MV-mediated g_H_+ decrease is dependent on AtPGR5.

We also investigated the effect of perturbing the thiol redox state of the cell on thylakoid conductivity in *Synechocystis*. As moderate concentrations of MV have only minor effects on cyanobacterial electron transport (Sétif, 2015), with no significant impact on g_H_+ observed after addition of 0.5 mM MV (Supplementary Fig. S15), we opted to test the effect of supplementation with the thiol-alkylating agent N-ethylmaleimide (NEM). Addition of 0.1 mM NEM to WT *Synechocystis* cultures resulted in a significant increase in g_H_+ after 5 seconds of illumination (Fig. 6a), similarly to results recently reported for another maleimide compound 2-methylmaleimide (Hubáček et al., 2024). In Δ*pgr5* cells, only a slight, statistically non-significant NEM-induced increase in g_H_+ was observed. In *Synechococcus*, which does not have PGR5, NEM did not have a significant effect on g_H_+ (Fig. 6b). Finally, we also tested the effect of NEM on g_H_+ in *Arabidopsis* WT and *pgr5* mutant leaves, repeating the fluctuating light experiment from Figure 1. NEM decreased g_H_+ throughout the experiment in WT (Fig. 6c), albeit not as dramatically as MV treatment. In contrast, *pgr5* mutant leaves did not respond to NEM treatment during the first LL-HL cycle (Fig. 6c). During subsequent cycles g_H_+ in NEM-treated *pgr5* mutant leaves decreased slightly in comparison to untreated *pgr5* mutant leaves, while still remaining significantly higher than untreated WT leaves (Supplementary Fig. S16) in HL.

**Figure 6.**
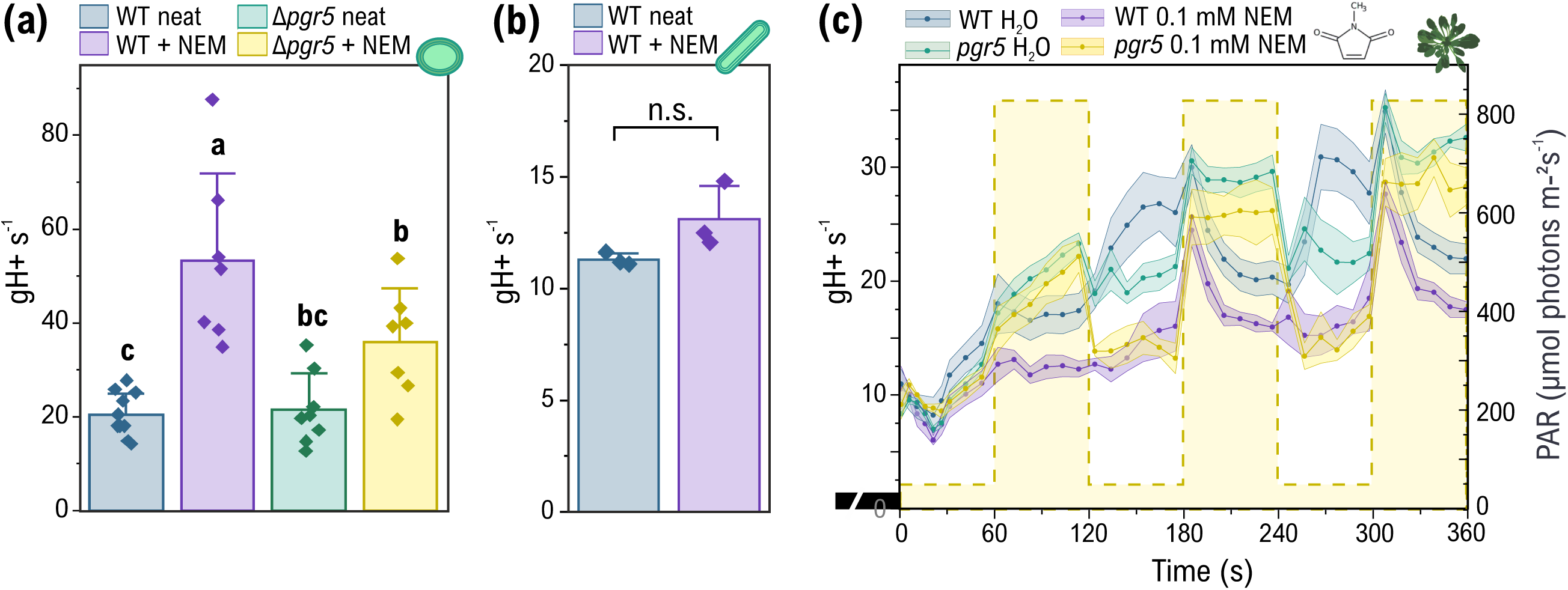
Effect of NEM on thylakoid conductivity under illumination. **(a)** The effect of 0.1 mM NEM on gH+ in *Synechocystis* WT and Δ*pgr5* cells. ECS DIRK were measured 5 seconds after the onset of illumination at 500 µmol photons m^-2^ s^-1^. The values are averages of 7–11 biological replicates + SD. Statistical significance of differences was tested by one-wayANOVA and Tukey’s tests for comparison of means (P<0.05**). (b)** The effect of 0.1 mM NEM on gH+ in *Synechococcus elongatus* sp. PCC 7942 WT cells. ECS DIRK were measured as averaged decay kinetics from 5 dark intervals spaced 5 seconds apart, with the first DIRK being administered 5 seconds after the onset of illumination at 500 µmol photons m^-2^ s^-1^. The values are averages of 3 biological replicates + SD. n.s. = statistically non-significant difference according to a paired *t*-test (P=0.20). **(c)** gH+ was measured in *Arabidopsis* WT-gl1 and *pgr5* mutant leaves incubated on water or water + 0.1 mM NEM. The water samples are the same as in Figure 1, while the +NEM values are averages of 4 biological replicates ± SEM.

These results provided evidence that PGR5 is involved in a redox-dependent regulatory mechanism of ATP synthase activity that is distinct from F_1_γ redox state. Interestingly, it has been reported that PGR5 from cucumber (*Cucumis sativus L.*) can interact with the AtpB subunit of the chloroplast ATP synthase in yeast-two-hybrid tests (Wu et al., 2021). To investigate the ability of PGR5 to directly interact with ATP synthase *in planta*, we performed bimolecular fluorescence complementation (BiFC) tests between AtPGR5 and AtCF_1_γ. Indeed, strong YFP fluorescence co-localising with chlorophyll autofluorescence was detected, indicating interaction between the two proteins in chloroplast thylakoid membranes (Fig. 7a). As negative controls we performed BiFC tests between AtPGR5 and the NdhS subunit of the chloroplast NAD(P)H dehydrogenase-like complex, another major thylakoid protein complex, as well as AtCF_1_γ and Thioredoxin x, a chloroplast protein similar in size to AtPGR5 (Supplementary Fig. S17). AtCF1γ +AtTRXf1 and AtPGRL1+AtPGR5 interactions were used as positive controls (Supplementary Fig. S18a–b), as both interactions are well established in previous studies (Nikkanen et al., 2016, 2018). No cleaved YFP fragments were detected in immunoblots with an antibody against YFP using in total protein extracts from *N. benthamiana* leaves transiently expressing the AtCF_1_γ:YFP-N fusion protein (Supplementary Fig. S18c).

**Figure 7.**
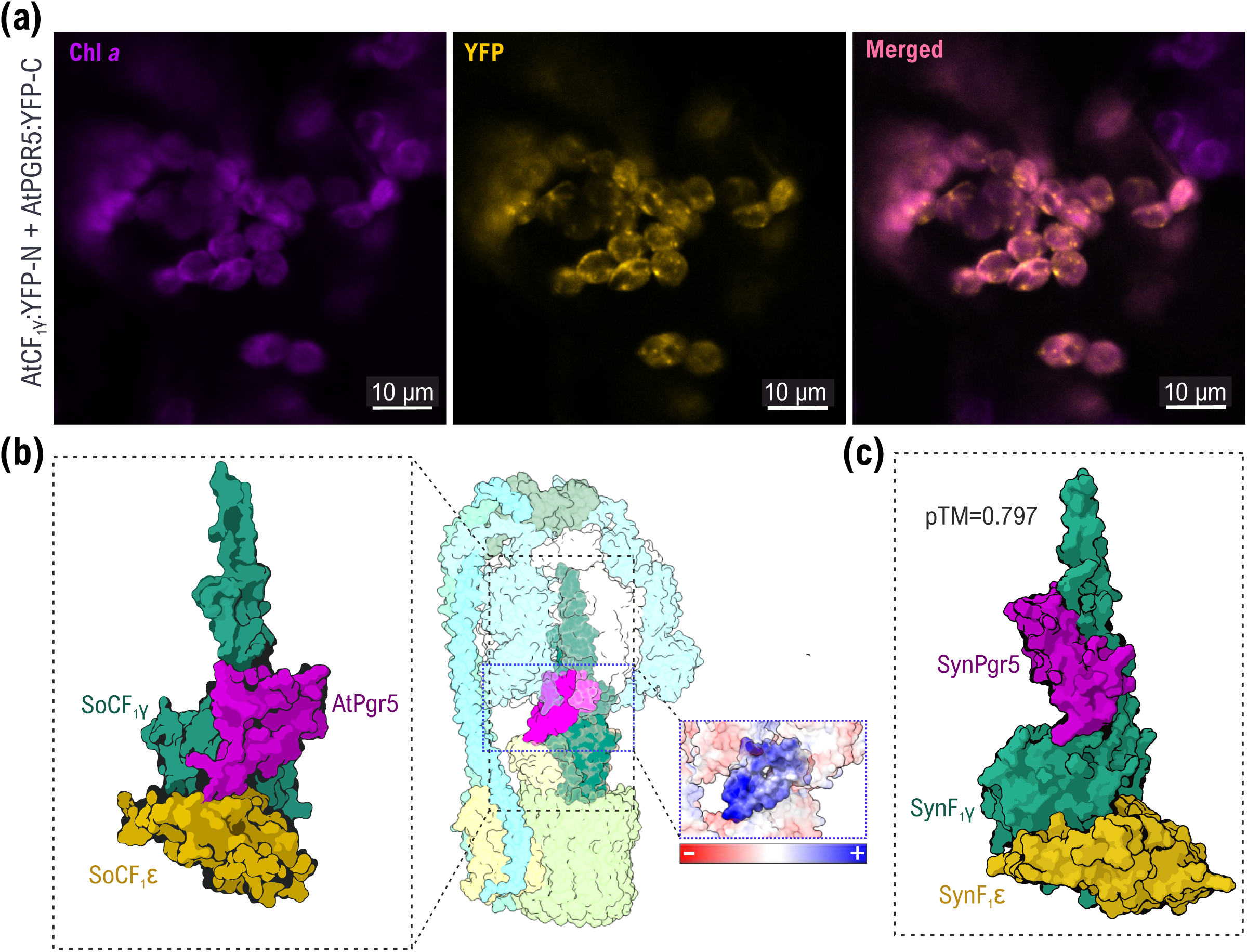
Protein–protein interaction between ATP synthase and PGR5. **(a)** Bimolecular fluorescence complementation (BiFC) test for protein-protein interaction between Arabidopsis CF_1_γ fused to an N-terminal fragment of yellow fluorescent protein (AtCF_1_γ:YFP-N) and Arabidopsis PGR5 fused to a C-terminal YFP fragment (AtPGR5:YFP-C) in tobacco leaves. The left panel shows Chlorophyll *a* autofluorescence in purple, the middle panel YFP fluorescence in yellow, and the right panel a merged image of YFP and chlorophyll images. The scale bar is 10 µm. **(b)** Predicted interaction complex of *Spinacia oleracea* CF_1_γ and CF_1_□ (Hahn et al., 2018), and AtPGR5 (PDB identifier Q9SL05) docked using the ClusPro 2.0 protein-protein docking service. The SoCF_1_γ / SoCF_1_□ / AtPGR5 with the of the Spinach ATP synthase complex (SoCF_0_CF_1_, PDB identifier 6FKF) superimposed (Hahn et al., 2018). Modelling of electrostatic surface charges of AtPGR5 and the surrounding parts of SoCF_1_ is shown in the smaller inlet on the right. **(c)** An Alphafold2 Multimer prediction of an interaction complex between *Synechocystis* ATP synthase γ subunit (*sll1327*, green), ATP synthase ε subunit (*slr1330,* yellow), and *Synechocystis* Pgr5 (*ssr2016*, magenta). The top-ranked model with a pTM score of 0.797 and an ipTM score of 0.702 is shown.

We also performed molecular docking of *Spinacia oleracea* CF_1_γ and CF_1_□ (Hahn et al., 2018) with an available AtPGR5 structural model using the ClusPro 2.0 protein-protein docking service (Kozakov et al., 2017). As the CF_1_ε-subunit likely remains bound to CF_1_γ even in its non-inhibitory conformation (Kato et al., 1997), we also included CF_1_ε in our model. The model predicts that PGR5 may slot between the CF_1_γ, CF_1_α, and CF_1_β subunits in a way that possibly hinders the rotation of CF_1_γ (Fig. 7b). Moreover, an Alphafold2 Multimer model predicts that SynPgr5 can bind to the F_1_γ subunit of the *Synechocystis* ATP synthase (Fig. 7c). We analysed the SoCF_1_γ / AtPGR5 (Supplementary Table S8) as well as SynF1γ / SynPgr5 (Supplementary Table S9) interaction interfaces using PDBePISA. Finally, we aligned the *Synechocystis* F_1_y sequences against the sequence from *Synechococcus elongatus* sp. PCC 7942, which does not have an PGR5 ortholog. Out of the residues mediating predicted hydrogen bonds with SynPgr5, only Pro 2 in the *Synechocystis* F1γ sequence is not conserved in the *Synechococcus* sequence (Supplementary Table S10). Therefore, further work is required to fully elucidate the molecular details of the mechanism of PGR5 interaction with ATP synthase.

## DISCUSSION

Substantial adjustments of thylakoid conductivity take place during sudden changes in light intensity in both chloroplasts and cyanobacteria, in order to control induction of the photoprotective mechanisms. Results shown herein indicate that this regulation is independent of the thiol redox state of CF_1_γ (Fig. 1), but dependent on another redox-based mechanism that involves the PGR5 protein in a manner that has been conserved in evolution from free-living cyanobacteria to green algae and C_3_ plants, as well as C_4_ plants.

### Does PGR5 have a role in CET?

The PGR5 protein was originally understood to either directly catalyse or facilitate CET around PSI in plants (Munekage et al., 2002) and cyanobacteria (Yeremenko et al., 2005). Lack of PGR5 caused a distinct inability to oxidise PSI in chloroplasts, which was interpreted as being caused by failure to induce photosynthetic control and NPQ due to insufficient acidification of the thylakoid lumen in the absence of PGR5-mediated CET (Yamamoto & Shikanai, 2019). However, contrasting evidence was already provided early on showing that ferredoxin-dependent CET does function in the absence of PGR5 (Nandha et al., 2007), suggesting that the phenotype of the *pgr5* mutant may be caused by another mechanism.

We showed previously that in *pgr5 Arabidopsis* plants overexpressing the chloroplast NADPH-dependent thioredoxin reductase (OE-NTRC *pgr5*), *pmf* was restored to a WT-level during LL-to-HL transitions (Nikkanen et al., 2018). This was insufficient however, to induce photosynthetic control, resulting in the OE-NTRC *pgr5* plants suffering from almost as severe over-reduction of the electron transport chain as *pgr5* plants. Concurring results were more recently reported from *Arabidopsis pgr5 cgl160* double mutants as well as plants expressing Plastid terminal oxidase 2 from *Chlamydomonas reinhardtii* (CrPTOX2) in the *pgr5* mutant background. Additional deletion of the ATP synthase assembly factor CGL160 resulted in a diminished amount of ATP synthase, limiting proton conductivity of the thylakoid membrane and allowing the *pgr5* mutant to maintain a WT-like *pmf* while still suffering from extreme acceptor side limitation of PSI (Kappel et al., 2023). Similarly, the introduction of CrPTOX2 in *pgr5* background resulted in restoration of WT-like (ΔpH-dependent) NPQ, but oxidation of PSI was still as impaired as in *pgr5* mutants (Zhou et al., 2023). These results suggest that the inability of *pgr5* mutants to induce photosynthetic control in HL is not simply caused by loss of ΔpH, but the AtPGR5 protein is directly required. It has been suggested that reduction of a disulphide in the Rieske subunit of Cyt b_6_f could upregulate photosynthetic control by causing a conformational change that allows protonation of the His128 residue at a higher lumenal pH (Hald et al., 2008; Degen & Johnson, 2024). Moreover, ΔpH-independent metabolic induction of photosynthetic control in *Chlamydomonas* was recently proposed by (Saroussi et al., 2023). It remains to be investigated how these potential mechanisms may be affected in PGR5-deficient lines, but as NTRC overexpression results in stronger reduction of chloroplast TRX systems and redox regulated proteins (Nikkanen et al., 2016), one could expect the activation of photosynthetic control in OE-NTRC *pgr5* to occur already at a lower ΔpH instead of being impaired.

Instead, the inability of *pgr5* mutants to oxidise PSI despite restoration of *pmf* generation may be explained by the presence of a partially photoinhibited PSI population with damaged F_A_ and F_B_ FeS clusters, resulting in repulsion of ferredoxin from the thylakoid membrane (Tiwari et al., 2024). This means that the bottleneck of electron transport in *pgr5* likely forms already at P700 or alternatively, as suggested by Buchert et al., (2020) for *Chlamydomonas*, at the Cyt b_6_f complex, and not at ferredoxin as would be expected if PGR5 was directly involved in ferredoxin-dependent CET. Accordingly, our near-infrared spectroscopy measurements showed that both reduction and re-oxidation kinetics of Fed are delayed in *pgr5* under far-red light while PC was more strongly oxidised (Supplementary Fig. S5). This indicated that the bottleneck is not in electron transfer onwards from Fed, but the accumulation of damaged PSI centres would make it appear as if the mechanism of ferredoxin-dependent CET is impaired. These results suggest that the phenotype of *pgr5* mutants may not be caused by direct impairment of CET, but it must be noted that we cannot fully exclude that possibility since impairment of CET and ATP synthase deregulation could both theoretically result in the original FeS cluster damage. Interestingly, the PSI oxidation phenotype of *pgr5* could be rescued via expression of *Physcomitrium patens* flavodiiron proteins in Arabidopsis *pgr5* background (Yamamoto et al., 2016), which likely provided a strong enough exogenous sink for electrons from ferredoxin to prevent PSI photodamage.

Finally, as it has been reported that a lack of CrPGR5 resulted in inefficient operation of the Q cycle (Buchert et al., 2020), it could be that the elevated gH+ values in *pgr5* mutants derive from diminished electrogenic activity of the Q cycle during the dark intervals where the rate of ECS signal decay is monitored. However, the post-illumination re-reduction rate of Cyt *f* was unimpaired in Arabidopsis *pgr5* (Supplementary Fig. S4). Moreover, Buchert et al. only detected a lower Cyt *f* re-reduction rate in anoxic *Chlamydomonas pgr5* samples. Therefore, it is unlikely that the elevated gH+ values in *pgr5* mutants are caused by impaired Q cycle activity.

The *Synechocystis* Δ*pgr5* mutant showed no impairment of CET (Fig. 2h). Our data is in line with a study by Miller et al. showing that the antimycin a-sensitive pathway (=PGR5-dependent) does not contribute to *pmf* formation during dark/light transitions in *Synechocystis*. Instead, the CET-dependent fraction of *pmf* generation was entirely attributable to the NDH-1 complex (Miller et al., 2021). As the extra ATP produced by CET is understood to enable more efficient operation of the CBC by adjusting the ATP:NADPH ratio, the fact that no impairment of CO_2_ fixation rate was observed in Δ*pgr5* (Supplementary Fig. S11e) further indicates that SynPgr5 is not needed for CET in *Synechocystis.* We propose it functions instead in reversible redox-dependent inhibition of ATP synthase.

Our ECS results both with the *Chlamydomonas pgr5* mutant (Fig. 3c) as well as the *pgr5-Cas Arabidopsis* mutant (Supplementary Fig. S6c) showed elevated proton flux (v_H_+) in HL, which is also incompatible with a significant role for PGR5 in CET, and may rather indicate enhancement of the CET rate. Accordingly, CrPGR5 was not detected in CET supercomplexes (Iwai et al., 2010).

Recruitment of FNR to thylakoid membranes was reported to be diminished in the absence of CrPGR5 (Mosebach et al., 2017), but a later study concluded that FNR remains associated with the thylakoids at all times (Kramer et al., 2021). It was also recently speculated, based on simultaneous near-infrared and chlorophyll fluorescence spectroscopy, that the absence of AtPGR5 may in fact enhance CET activity in *Arabidopsis* (Maekawa et al., 2024). A lower v_H_+ in LL that cannot be fully explained by a lower linear electron transport rate has, however, been reported in *Arabidopsis* (Degen et al., 2023) and *S. viridis* (Woodford et al., 2024) *pgr5* mutants. Interestingly though, the *Arabidopsis pgr5-Cas* mutant showed WT-like v_H_+ in LL (Supplementary Fig. S6c), suggesting that the low v_H_+ in the *pgr5-1* mutant (Fig. 1c) may be affected by the additional mutation of the At2g1724 gene (Wada et al., 2021). Moreover, in the fluctuating light conditions of the current study, no impairment of vH+ was detected in the *S. viridis pgr5* mutant (Supplementary Fig. S14c). It should be noted, however, that the presence of damaged PSI centres in *pgr5* mutants would result in a decrease in the ECS signal induced by a single-turnover flash (ECS_ST_), which was used to normalise the ECS values in plant and algal samples. If this decrease in ECS_ST_ is not proportional to a decrease in the actinic light-induced ECS signal, it may result in overestimation of the *pmf* and vH+ parameters. However, normalisation of the *Chlamydomonas* ECS data (Fig. 3) to [Chl] only did not eliminate the differences between WT and *pgr5* (Supplementary Fig. S13), suggesting that no major artefact is caused by potential PSI damage in *pgr5*.

### Conserved PGR5-mediated regulatory mechanism of ATP synthase depends on thiol redox conditions

The g_H_+ values elevated independently of *pmf* in PGR5-deficient *Synechocystis*, *Chlamydomonas*, *Arabidopsis,* and *S. viridis* during high-light phases of fluctuating light (Fig. 1–4, Supplementary Fig. S14) are consistent with PGR5 functioning as a direct or indirect down-regulator of ATP synthase activity (Tikkanen et al., 2015; Kanazawa et al., 2017; Nikkanen et al., 2018). *In planta* interaction of AtPGR5 with AtCF_1_γ in BiFC tests (Fig. 7a) and *in silico* modelling of PGR5-CF_1_γ interaction complexes (Fig. 7b–c) suggest that PGR5 may inhibit ATP synthase by directly binding to the F_1_γ subunit. F_1_ε-inhibition likely prevents ATP hydrolysis in dark and LL conditions, when *pmf* is low, while PGR5 would limit ATP synthase activity in HL, possibly allowed by the non-inhibitory “down” conformation of F_1_ε (Feniouk et al., 2010), to allow build-up of sufficient ΔpH for induction of photoprotection.

The mechanism PGR5-dependent inhibition of ATP synthase activity may be regulated by the thiol redox state of the chloroplast stroma or cell. PGR5 contains a conserved cysteine residue (Munekage et al., 2002; Yeremenko et al., 2005), which could be target for regulatory thiol modifications. In line with this hypothesis, AtPGR5 was found able to interact with AtNTRC in an *in planta* BiFC test, and overexpression of AtNTRC resulted in intensified downregulation of thylakoid proton conductivity during transitions to HL, but only in the presence of AtPGR5 (Nikkanen et al., 2018). Moreover, using a *pgr5 ntrc* double mutant of *Arabidopsis*, Naranjo et al. showed that the high NPQ observed in *ntrc* mutant plants is dependent on the presence of AtPGR5 (Naranjo et al., 2021). *Ntrc* mutant plants have a diminished AtPGR5 protein level (Nikkanen et al., 2018), most likely to compensate for the impaired activation of ATP synthase via reduction of the AtCF_1_γ disulphide (Carrillo et al., 2016; Nikkanen et al., 2016).

The *pgr5* mutation could, hypothetically, result in the chloroplast thioredoxin systems themselves becoming more oxidised or reduced, which would affect the redox state and activity of some other inhibitory factor of ATP synthase. However, no differences were reported between Arabidopsis WT and *pgr5* in the vivo redox states of redox regulated CBC enzymes FBPase and SBPase, of NADP-malate dehydrogenase, or of CF_1_γ (Okegawa & Motohashi, 2020) (Fig 1g). These findings indicate that altered activity of the thioredoxin systems is not a major factor in the gH+ phenotype of *pgr5*. A small population of reduced CF_1_γ was detected in *pgr5* in the dark (Fig. 1g), but this would not affect the gH+ dynamics under fluctuating light apart from the very first seconds of the dark-to-light transition.

It has been suggested that a disulphide bridge between AtPGR5 and AtPGRL1 is required for PGR5/PGRL1-dependent CET in *Arabidopsis* (Hertle et al., 2013). More recently, however, a model was proposed where the function of AtPGRL1 is to stabilise AtPGR5 (Rühle *et al.,* 2021). Moreover, *Chlamydomonas* PGR5 with the conserved cysteine residue mutated to a serine was able to complement the impairment of CET in the *Chlamydomonas pgr5* mutant (Buchert et al., 2020), suggesting that the conserved cysteine residue in PGR5 may be involved in a non-CET related regulatory mechanism.

Transient reduction of AtPGRL1 occurs upon onset of illumination, coinciding with increase in *pmf*, PSI oxidation, and induction of NPQ (Hertle et al., 2013; Nikkanen et al., 2018; Wolf et al., 2020; Chaturvedi et al., 2024). TRXm4 forms a disulphide bridge with AtPGRL1, and cleavage of that disulphide (most likely by another thioredoxin isoform) and transient release of TRXm4 also occur at dark-to-light transitions (Okegawa & Motohashi, 2020). This has been suggested to indicate activation of CET (Hertle et al., 2013; Wolf et al., 2020; Okegawa & Motohashi, 2020; Chaturvedi et al., 2024), but the same results could also be interpreted as reduction of AtPGRL1 affecting the association of AtPGR5 with AtPGRL1, releasing AtPGR5 to interact with ATP synthase and to downregulate its activity (Fig. 8). This would limit H^+^ efflux from the lumen and cause build-up of *pmf,* and consequently, induction of NPQ and photosynthetic control. As recently proposed for *S. viridis* (Nix et al., 2024), PGR5 may be bound to PGRL1 homodimers that are held together by an intermolecular disulphide between the Cys 116 residues of the monomers, possibly maintained by a bound TRXm4. Once stromal redox regulatory systems become sufficiently reduced, the disulphide would be reduced, potentially by another TRX isoform, dissolving the PGRL1 dimer and releasing PGR5.

**Figure 8.**
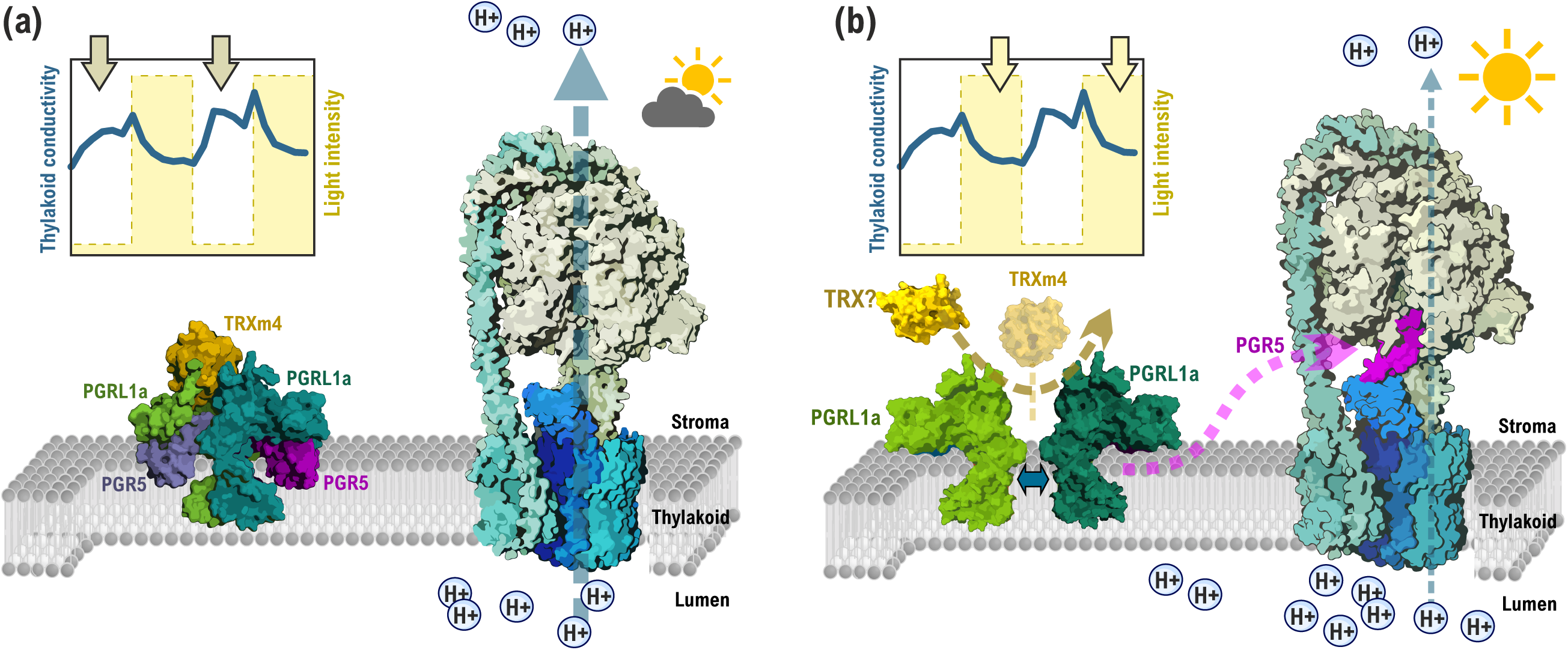
**Hypothetical model of PGR5-mediated downregulation of ATP synthase activity in plant chloroplasts**. A model of an *Arabidopsis thaliana* 2xPGRL1a/2xPGR5/TRXm4 complex was created with the protein–protein docking service ClusPro 2.0, docking monomeric AlphaFold models with the PDB identifiers Q8H112, Q9SL05, and Q9SEU6. An interaction complex of AtPGR5, AtCF_1_γ, and AtCF_1_□ was modelled with AlphaFold3 and overlaid with the from the *Spinacia oleracea* CF_0_CF_1_ crystal structure (6FKF). **(a)** Under low irradiance when stromal redox regulatory systems are largely oxidised, PGR5 is bound to the PGRL1a/PGR5/TRXm4 complex, possibly held by an intermolecular disulphide bond between the Cys116 residues in the PGRL1a monomers. This allows ATP synthase to maintain high activity, effectively dissipating the proton gradient over the thylakoid membrane**. (b)** Under high irradiance, stromal redox regulatory systems become reduced, resulting in reduction of the PGRL1a-dimer disulphide and release of PGR5 to bind to ATP synthase. This inhibits the activity of ATP synthase, lowering thylakoid conductivity to protons, buildup of high *pmf* and induction of photoprotective mechanisms.

We therefore suggest that the phenotypes of *pgr5* mutants may have been misinterpreted as being caused by directly impaired CET. For example, it was recently shown that point mutations of disulphide-forming cysteine residues in AtPGRL1a results in elevated *pmf*, NPQ, and donor-side limitation of PSI (Chaturvedi et al., 2024). While the authors did not show data on g_H_+, it is possible that this phenotype was caused not by increased CET, but by constitutive inhibition of ATP synthase activity by AtPGR5. Similarly, it has been reported that AtPGR5 can enhance CET if both AtPGRL1 and AtPGRL2, a protein causing degradation of AtPGR5 in the absence of AtPGRL1, are missing (Rühle et al., 2021). This conclusion was based on higher *pmf* and NPQ in the *pgrl1a pgrl1b pgrl2* triple mutant compared to *pgrl1* or *pgr5* single mutants. We suggest, however, that the increased *pmf* was, again, probably mediated or at least contributed to by lowered ATP synthase conductivity.

In the current study, we explored the effects of perturbing the chloroplast or cellular redox state on the conductivity of ATP synthase. Interestingly, we found that addition of methyl viologen (MV) results, either directly or e.g. via affecting the TrxL2-dependent oxidative pathway (Yoshida et al., 2018), in strong reduction of chloroplast enzymes in *Arabidopsis*, including AtCF_1_γ (Fig. 5a). While we cannot fully exclude the possibility of the apparent reduction of chloroplast enzymes in presence of MV being caused by overoxidation of thiols to sulphenic acids (Huang et al., 2019), the elevation of g_H_+ at the onset of illumination in MV-treated leaves (Fig. 5b) suggests otherwise, at least in the case of CF_1_γ in darkness. Moreover, overoxidation would inactivate 2-CysPrxs, preventing their oxidising effect on other thiol proteins in the chloroplast, (Pérez-Ruiz et al., 2017; Vaseghi et al., 2018). Thus, we conclude that MV treatment indeed results in strong reduction of chloroplast redox enzymes, including AtCF_1_γ.

Accordingly, g_H_+ was elevated in dark-adapted MV-treated leaves (Fig. 5b). However, strong downregulation of g_H_+ during illumination was measured in MV-leaves, but only in the presence of AtPGR5 (Fig. 5c). Thiol-alkylating agent NEM had a similar albeit less dramatic effect (Fig. 6c).

Interestingly, lack of AtPGR5 made the plants immune to NEM-induced g_H_+ decrease under HL, but did not prevent the decrease observed under LL phases of the fluctuating light experiment. These results provide further evidence that AtPGR5 is required for redox-dependent downregulation of g_H_+ under high irradiance, although it should be noted that we cannot exclude indirect effects of MV/NEM treatment interfering with metabolic regulation of ATP synthase (Kohzuma et al., 2013). We hypothesise that in WT *Arabidopsis*, addition of MV likely causes strong reduction of either AtPGR5/AtPGRL1/AtTRXm4 directly or some associated factor that would, without an MV treatment, function as an inhibitor of the ATP synthase during sudden increases in light intensity. Importantly, this inhibitory effect overrides the activating effect of a fully reduced AtCF_1_γ. In the absence of AtPGR5, the inhibitory mechanism does not function even in presence of MV.

In *Synechocystis*, addition of NEM caused an opposite phenotype to *Arabidopsis*, with g_H_+ being elevated rather than diminished (Fig. 6a). The NEM-induced g_H_+ increase was diminished in *Synechocystis* Δ*pgr5* and absent in WT *Synechococcus*, a species that does not have a PGR5 orthologue (Shimakawa et al., 2016). Untreated *Synechococcus* did, however, exhibit similar downregulation of g_H_+ under strong illumination as *Synechocystis* or *Arabidopsis* (Supplementary Fig. S12). These results suggest that the regulatory mechanisms involved in this response may differ between organisms in terms of directionality of redox-regulation and involvement of PGR5, with *Synechococcus* possibly using an unrelated, convergently evolved analogous mechanism. The molecular details of these differing responses remain to be elucidated in future studies.

Interestingly, it has been reported that alkylation of cysteine 89 (Cys 130 in the full sequence containing the transit peptide) in the (*Spinacia oleracea*) SoCF_1_γ by NEM or crosslinking Cys89 with Cys322 inhibits photophosphorylation and induces loss of the proton gradient (McCarty & Fagan, 1973; Soteropoulos et al., 1994; Evron & Pick, 1997). This effect was not observed in the dark, suggesting Cys89 only becomes accessible to modifications in light upon generation of ΔpH (McCarty & Fagan, 1973). It is therefore possible that Cys89 is responsible for the middle band in the AtCF_1_γ mobility shift assays that is also present in the control sample (Fig. 1e), where all thiols accessible in the dark are alkylated with NEM before addition of MAL-PEG. MAL-PEG likely gains access to Cys89 after the subsequent DTT treatment, resulting in the observed mobility shift. The F_1_γ Cys89 residue is conserved in the green lineage, including cyanobacteria (Hisabori et al., 2013), suggesting a functional role. Our computational models of AtCF_1_γ-AtPGR5 and SynF_1_γ-SynPgr5 interaction complexes predict, however, that neither the conserved Cys30 in SynPgr5, or Cys90 in SynF_1_γ are involved in mediating the protein–protein interactions (Supplementary Tables S1, S2). This is also consistent with the lack of changes in AtCF_1_γ redox state in fluctuating light (Fig. 1f–g), suggesting that thiol/disulphide exchanges of F_1_γ are not involved in the mechanism of adjusting g_H_+ in fluctuating light. The predicted binding of SynPgr5 to SynF_1_γ in our model somewhat resembles the binding of the inhibitory IF_1_ protein to mitochondrial ATP synthase, which slots between the F_1_γ, F_1_α, and F_1_β subunits (García-Bermúdez & Cuezva, 2016). This could also explain the observed interaction between cucumber PGR5 and F_1_β (Wu et al., 2021).

An alternative or additional mechanism for the regulation of g_H_+ in fluctuating light in plants could function through changes in intercellular inorganic carbon concentration (C_i_). C_i_ increases upon HL to LL transitions as carbon fixation rate suddenly drops (Vialet-Chabrand et al., 2017), and increase in C_i_ is known to increase g_H_+ via an unknown regulatory pathway (Kohzuma et al., 2013). It is unlikely, however, that the inability of *pgr5* to down-regulate g_H_+ in HL would be caused by an elevated concentration of C_i_, because WT-like levels of C_i_ as well as CO_2_ fixation rates were measured under HL in PGR5-deficient rice plants (Nishikawa et al., 2012; Wada et al., 2018). The carbon fixation capacity of *pgr5* plants only goes down as a consequence of prolonged HL exposure and ensuing PSI photoinhibition (Gollan et al., 2017; Lima-Melo et al., 2018). Similarly, we detected no impairment of CO_2_ fixation in the *Synechocystis* Δ*pgr5* mutant (Supplementary Fig. S11e). Thus, it is also unlikely that the elevated gH+ in PGR5-deficient plants or cyanobacteria would be caused by an excess of P_i_ due to a hyperactivated CBC. Interestingly, expression of FlvA and FlvB from the moss *Physcomitrium patens* in the Arabidopsis *pgr5* mutant background alleviated the elevated g_H_+ phenotype (Yamamoto & Shikanai, 2020), highlighting the complex relationship between oxidation of PSI and regulation of ATP synthase.

### Physiological significance of PGR5-mediated downregulation of ATP synthase in *Synechocystis*

Lack of SynPgr5 caused no impairment of PSI oxidation or growth despite lowered *pmf* in *Synechocystis* under fluctuating light (Fig. 2). This may be due to prevalence of FDPs as a protective electron sink on the acceptor side of PSI in cyanobacteria (Allahverdiyeva et al., 2013; Santana-Sanchez et al., 2019). Slightly elevated FDP activity was indeed measured in fluctuating light grown Δ*pgr5* under fluctuating light (Fig. 2i), which may allow the mutant to maintain sufficient PSI oxidation in HL despite diminished *pmf* and to grow similarly to WT in fluctuating light (Fig. 2g). We have recently proposed that dissipation of *pmf* (lower cytosolic pH) causes FDP hetero-oligomers to associate more strongly with the thylakoid membrane, increasing photoprotective O_2_ photoreduction activity (Eckardt et al., 2024; Nikkanen et al., 2025). In Δ*pgr5,* the decreased *pmf* due to elevated g_H_+ in HL (Fig. 2a–b) may therefore cause the slightly increased FDP activity during the HL phases of fluctuating light conditions (Fig. 2i), at least partly compensating for the loss of photoprotective capacity via ΔpH-induced photosynthetic control. Even in the absence of Flv1/3-mediated O_2_ photoreduction in the Δ*pgr5 flv3* double mutant, however, no significant additional impairment of PSI oxidation in comparison to Δ*flv3* single mutant was observed (Fig. 2e–f). The lack of any clear effect on PSI oxidation in Δ*pgr5* is thus most likely explained by PSI/PSII stoichiometry, as *Synechocystis* harbours 2-to-5-fold more PSI than PSII complexes (Moore & Vermaas, 2024).

Interestingly, there appears to be a regulatory link between FDPs, ATP synthase, and SynPgr5 in cyanobacteria. During transitions to HL, lack of Flv1/3 results in a loss up to 75% of the proton flux during the first minute of illumination in *Synechocystis* (Nikkanen et al., 2020) (Fig. 2c) as well as in the filamentous model cyanobacterium *Anabaena* sp. PCC 7120 (Santana-Sánchez et al., 2023).

However, cyanobacterial cells compensate for this by downregulating g_H_+ (Fig. 2a), achieving WT-like *pmf* levels after the initial stages of HL transitions (Fig. 2b). In in the absence of both Flv1/3 and SynPgr5 in the Δ*pgr5 flv3* double mutant, however, this downregulation of g_H_+ did not occur, indicating that SynPgr5-mediated inhibition of ATP-synthase is required for the compensatory mechanism. Interestingly, proton flux also recovered to WT-levels in Δ*pgr5 flv3* (Fig. 2c), suggesting e.g. increased CET via NDH-1.

It remains to be investigated what the environmental conditions are where SynPgr5 has a physiologically significant role. Future studies should focus for example on mixotrophic conditions, where overexpression of SynPgr5 has been shown to inhibit growth (Margulis et al., 2020) and diurnal rhythms, which present additional challenges for coordination of the bioenergetic pathways and metabolism in cyanobacteria (Solymosi et al., 2020; Ortega-Martínez et al., 2024).

## Conclusion

The activity of chloroplast or cyanobacterial ATP synthase is dynamically regulated during fluctuations in light intensity. These adjustments are not dependent on the thiol redox state of the F_1_γ subunit, but instead depend on the PGR5 protein as well as the thiol redox state of the stroma or cytosol. This mechanism has been conserved in evolution from free-living cyanobacteria to chloroplasts in algae, C_3_ plants as well as C_4_ plants. The inability to downregulate ATP synthase activity under strong illumination is the only phenotype of PGR5-deficient knockout mutants that can be observed in all representatives of the groups of photosynthetic organisms studied here; cyanobacterium *Synechocystis* sp. PCC 6803, green alga *Chlamydomonas reinhardtii*, C_3_ plant *Arabidopsis thaliana*, and C_4_ plant *Setaria viridis*. The role of PGR5 is vital for oxidation of PSI in plants and algae, but not in cyanobacteria. We also demonstrated that the cyanobacterial Pgr5 is not needed for cyclic electron transport. We suggest that PGR5 directly binds to ATP synthase and functions as an inhibitory factor in a manner that is dependent on the thiol redox state of the cell or chloroplast. The observed effect of plant PGR5 on CET is likely an indirect consequence of misregulation of ATP synthase and, as shown by (Tiwari et al., 2024), increased photoinhibition of PSI, although a role for PGR5 in regulating CET via Cyt b_6_f cannot be excluded (Buchert et al., 2020). Our study lays important groundwork for understanding the regulatory mechanisms of the *pmf* and their evolution, possibly enabling biotechnologies to modulate *pmf* regulation for enhancement of photosynthetic efficiency in crop plants or microbial bioproduction platforms.

## Supplementary Data

**Fig. S1.** PCR analysis to verify full segregation of the Δ*pgr5* and Δ*pgr5 flv3* strains.,

**Fig. S2**. Mobility shift assay of the in vivo thiol redox state of CBC enzyme fructose 1,6 bisphosphatase (FBPase).

**Fig. S3.** PSI quantum yield in PGR5-mutants of Arabidopsis and *Synechocystis*.

**Fig. S4.** Cytochrome *f* post-illumination re-reduction rates in Arabidopsis WT and *pgr5-1* mutant leaves.

**Fig. S5**. Redox kinetics of PC, P700, and Fed upon a pulse of far-red light in Arabidopsis WT and *pgr5*.

**Fig. S6.** *Pmf* dynamics in the *Arabidopsis thaliana* WT and *pgr5-Cas* mutant in fluctuating light.

**Fig. S7.** *Pmf* dynamics in the *Arabidopsis thaliana* WT and *pgr5*-Cas mutant in fluctuating light in the presence of inhibitors of mitochondrial respiration

**Fig. S8.** Immunodetection of AtpB and Flv3 levels in *Synechocystis* strains.

**Fig. S9.** *In vivo* redox kinetics of P700, PC, and Fed and O_2_ and CO_2_ gas exchange in Δ*pgr5 flv3* and Δ*flv3* strains of *Synechocystis*.

**Fig. S10.** P700 oxidation under far-red light in WT and Δ*pgr5* strains of *Synechocystis*.

**Fig. S11.** Photosynthetic phenotype of the *Synechocystis* Δ*pgr5* strain.

**Fig. S12.** *Pmf* dynamics in *Synechococcus elongatus* sp. PCC 7942 under fluctuating light.

**Fig, S13.** *Pmf* and vH+ dynamics in *Chlamydomonas reinhardtii* under fluctuating light without ECS_ST_ normalisation.

**Fig. S14.** *Pmf* dynamics in the C_4_ model grass *Setaria viridis*.

**Fig. S15.** Effect of methyl viologen on thylakoid conductivity in *Synechocystis*.

**Fig. S16.** Effect of N-ethylmaleimide (NEM) on thylakoid conductivity under fluctuating light in Arabidopsis.

**Fig. S17.** Negative controls for the BiFC tests.

**Fig. S18.** Positive controls for bimolecular fluorescence com plementation (BiFC) tests.

**Table S1.** LMEM report table for Arabidopsis WT vs *pgr5-1* (full data).

**Table S2.** LMEM report table for Arabidopsis WT vs *pgr5-1* (high light data)

**Table S3.** LMEM report table for Arabidopsis WT vs *pgr5-*Cas (full data).

**Table S4.** LMEM report table for Arabidopsis WT vs *pgr5-*Cas (high light only).

**Table S5.** LMEM report table for *Chlamydomonas* WT vs *pgr5* (full data).

**Table S6.** LMEM report table for *Setaria viridis* WT vs *pgr5-1* (full data).

**Table S7**. LMEM report table for *Synechocystis* WT vs Δ*pgr5* (full data).

**Table S8.** Hydrogen Bond Interactions and salt bridges between AtPGR5 and SoCF_1_γ in the ClusPro 2.0 docking model.

**Table S9.** Hydrogen Bond Interactions between SynCF_1_γ and SynPgr5 in the AlphaFold2 model.

**Table S10**. Alignment of the ATP synthase F1γ amino acid sequences from Synechocystis sp. PCC 6803 and Synechococcus elongatus sp. PCC 7942

## Acknowledgements

We thank Drs Martina Jokel and Arjun Tiwari for providing *Chlamydomonas* strains and the Arabidopsis *pgr5*-Cas line, respectively. Linda Nevala, Mahfuzur Rahman, Iida Turpeinen, and Tapio Ronkainen are thanked for technical assistance, and the Turku Bioscience CIC Imaging Core for imaging infrastructure and assistance.

## Author Contributions

L.N. conceived the research idea, devised the experimental plan, performed most of the experiments, analysed the data, and wrote the first draft of the article. L.T.W. performed part of the cyanobacterial ECS and Dual-KLAS-NIR measurements and analysed the data. R.W. and M.E. performed the ECS measurements in *S. viridis*. H.M. created the *Synechocystis* Δ*pgr5* and Δ*pgr5 flv3* mutants. D.K. performed *Chlamydomonas* cell culture and ECS measurements. E.R. and Y.A. supervised research and provided resources. All authors participated in editing the article.

## Data Availability

The SynPgr5/SynCF1γ AlphaFold2 multimer model is available in the ModelArchive database at https://www.modelarchive.org/doi/10.5452/ma-v9v6q. All relevant data generated in this study are included in this paper and its Supplementary Data.

