## Supplementary Data for "PGR5 is needed for redox-dependent regulation of ATP synthase both in chloroplasts and in cyanobacteria"

The following Supplementary Figures and Tables are available for this article:

**Fig. S1.** PCR analysis to verify full segregation of the Δ*pgr5* and Δ*pgr5 flv3* strains.,

**Fig. S2**. Mobility shift assay of the in vivo thiol redox state of CBC enzyme fructose 1,6 bisphosphatase (FBPase).

**Fig. S3.** PSI quantum yield in PGR5-mutants of Arabidopsis and *Synechocystis*.

**Fig. S4.** Cytochrome *f* post-illumination re-reduction rates in Arabidopsis WT and *pgr5-1* mutant leaves.

**Fig. S5**. Redox kinetics of PC, P700, and Fed upon a pulse of far-red light in Arabidopsis WT and *pgr5*.

**Fig. S6.** *Pmf* dynamics in the *Arabidopsis thaliana* WT and *pgr5-Cas* mutant in fluctuating light.

**Fig. S7*.*** *Pmf* dynamics in the *Arabidopsis* *thaliana* WT and *pgr5*-Cas mutant in fluctuating light in the presence of inhibitors of mitochondrial respiration

**Fig. S8.** Immunodetection of AtpB and Flv3 levels in *Synechocystis* strains.

**Fig. S9.** *In vivo* redox kinetics of P700, PC, and Fed and O_2_ and CO_2_ gas exchange in *Δpgr5* *flv3* and Δ*flv3* strains of *Synechocystis*.

**Fig. S10.** P700 oxidation under far-red light in WT and Δ*pgr5* strains of *Synechocystis*.

**Fig. S11.** Photosynthetic phenotype of the *Synechocystis Δpgr5* strain.

**Fig. S12.** *Pmf* dynamics in *Synechococcus elongatus* sp. PCC 7942 under fluctuating light.

**Fig, S13.** *Pmf* and vH+ dynamics in *Chlamydomonas reinhardtii* under fluctuating light without ECS_ST_ normalisation.

**Fig. S14.** *Pmf* dynamics in the C_4_ model grass *Setaria viridis.*

**Fig. S15.** Effect of methyl viologen on thylakoid conductivity in *Synechocystis*.

**Fig. S16.** Effect of N-ethylmaleimide (NEM) on thylakoid conductivity under fluctuating light in Arabidopsis.

**Fig. S17.** Negative controls for the BiFC tests.

**Fig. S18.** Positive controls for bimolecular fluorescence com

plementation (BiFC) tests.

**Table S1.** LMEM report table for Arabidopsis WT vs *pgr5-1* (full data)*.*

**Table S2.** LMEM report table for Arabidopsis WT vs *pgr5-1* (high light data)*.*

**Table S3.** LMEM report table for Arabidopsis WT vs *pgr5-*Cas (full data)*.*

**Table S4.** LMEM report table for Arabidopsis WT vs *pgr5-*Cas (high light only)*.*

**Table S5.** LMEM report table for *Chlamydomonas* WT vs *pgr5* (full data)*.*

**Table S6.** LMEM report table for *Setaria viridis* WT vs *pgr5-1* (full data)*.*

**Table S7**. LMEM report table for *Synechocystis* WT vs Δ*pgr5* (full data)*.*

**Table S8.** Hydrogen Bond Interactions and salt bridges between AtPGR5 and SoCF_1_γ in the ClusPro 2.0 docking model.

**Table S9.** Hydrogen Bond Interactions between SynCF_1_γ and SynPgr5 in the AlphaFold2 model.

**Table S10**. Alignment of the ATP synthase F1γ amino acid sequences from Synechocystis sp. PCC 6803 and Synechococcus elongatus sp. PCC 7942

**Supplementary Figure S1.** PCR analysis from genomic DNA to verify full segregation of the Δ*pgr5* and Δ*pgr5 flv3* strains.

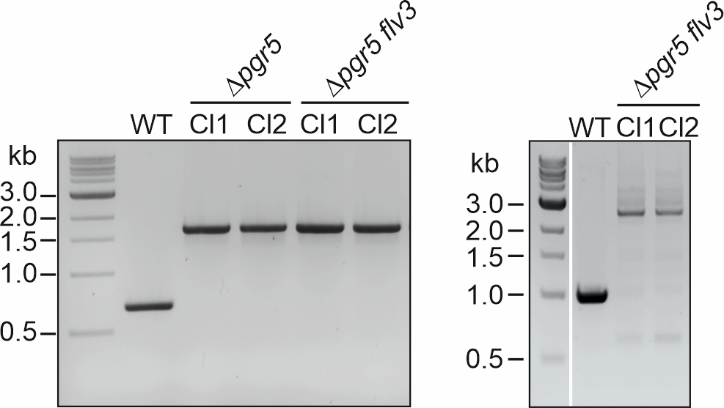

**Supplementary Figure S2**. **Mobility shift assay of the in vivo thiol redox state of the Calvin–Benson cycle enzyme fructose 1,6 bisphosphatase (FBPase)**. Samples of WT Arabidopsis leaves were taken from fluctuating light (1 min periods of 50 µmol photons m^-2^s^-1^, LL, and 825 µmol photons m^-2^s^-1^, HL.

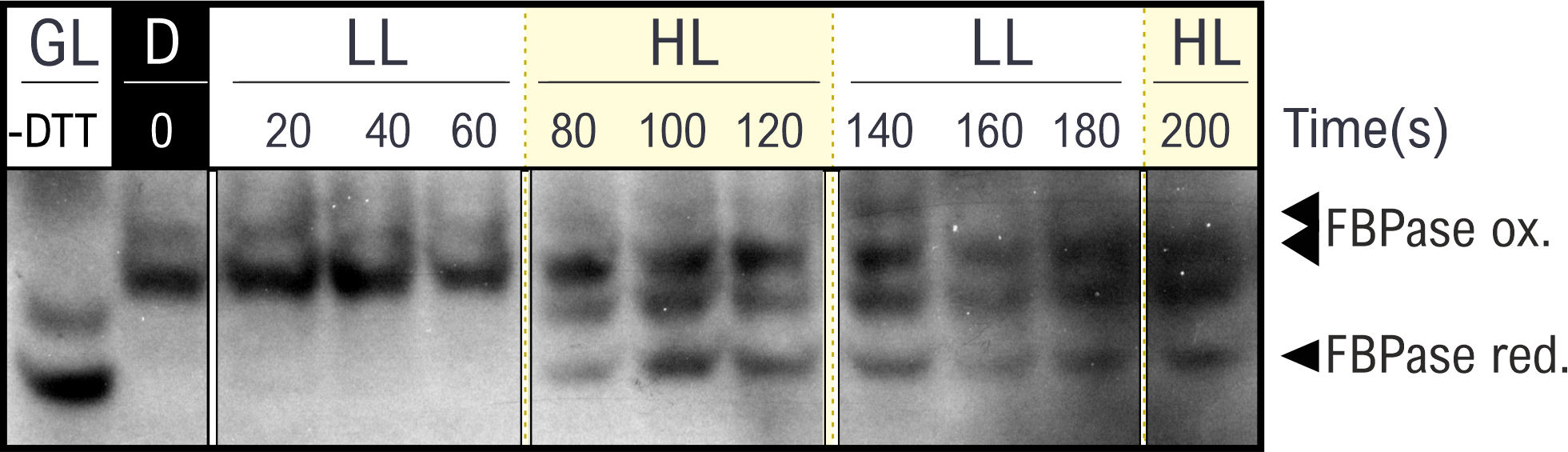

**Supplementary Figure S3. PSI quantum yield in PGR5-mutants of Arabidopsis and *Synechocystis*.**

**(a)** PSI quantum yield (Y(I)) and **(b)** PSI acceptor side limitation (Y(NA)) in Arabidopsis WT and *pgr5-1* mutant leaves in fluctuating light, as measured by the Dual-PAM 100 spectrophotometer. Values in a–b are averages from 3 (WT) and 5 (pgr5-1) biological replicates ±SEM, with statistically significant differences according to two-tailed Student’s t-tests indicated by asterisks. **(c)** Y(I) in *Synechocystis* WT and Δ*pgr5,* Δ*pgr5 flv3* and Δ*flv3* mutant cells in fluctuating light. Values are averages from 3–6 biological replicates ±SEM, with statistically significant differences between WT and *Δpgr5* according to two-tailed Student’s t-tests indicated by asterisks.

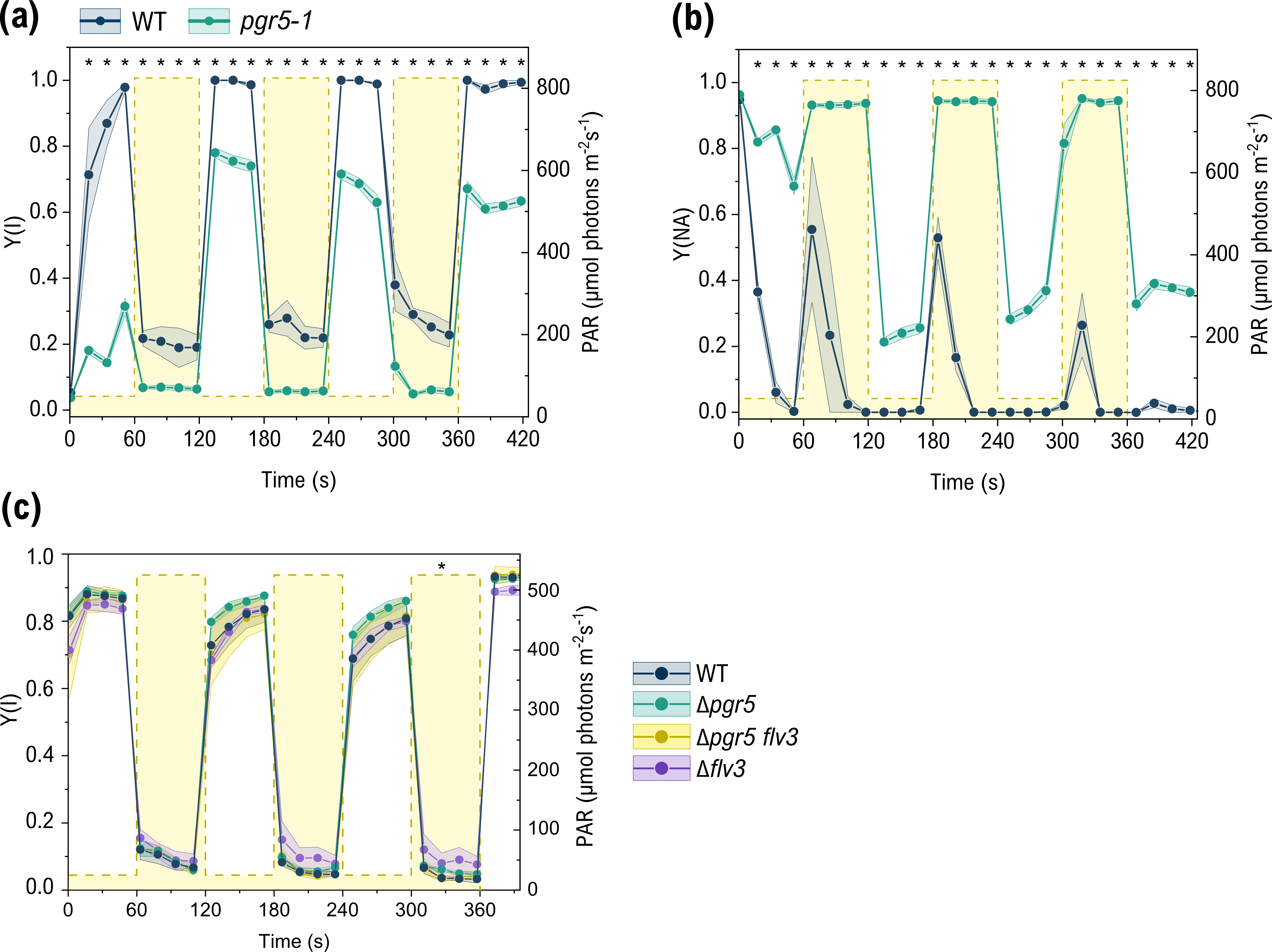

**Supplementary Figure S4. Cytochrome *f* post-illumination re-reduction rates in Arabidopsis WT and *pgr5-1* mutant leaves.** Cytochrome *f* redox kinetics were measured as time-resolved absorbance changes at 554 nm with a baseline drawn between 546 and 573 nm during and after five seconds of illumination with 500 µmol photons m^-2^ s^-1^ of green light, using the JTS-10 spectrophotometer. The cytochrome f post-illumination re-reduction rates (*k_f_* _red_ s^-1^) were obtained by fitting the post-illumination kinetics over 300 ms to a first-order exponential function. Left panel: average *k_f_* _red_ s^-1^ values from four biological replicates ± SEM is shown, with individual data points as dots. No statistically significant difference was detected between WT and *pgr5* according to a Student’s t-test.

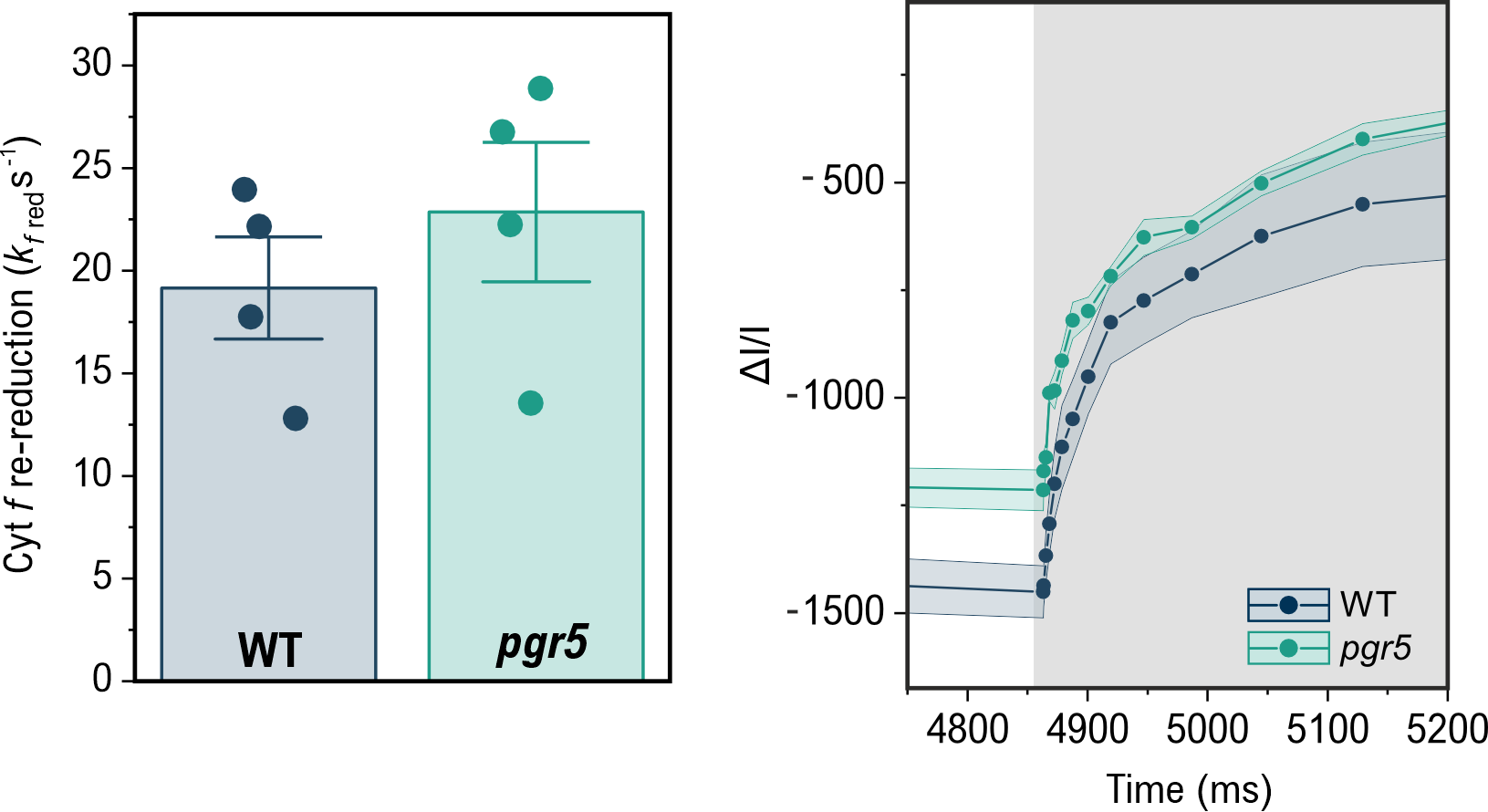

**Supplementary Figure S5. Redox kinetics of PC, P700, and Fed upon a pulse of far-red light in Arabidopsis WT and *pgr5*.** The redox kinetics of plastocyanin (PC), P700 and ferredoxin (Fed) were deconvoluted from near-infrared absorbance difference measurements using the Dual-KLAS-NIR spectrophotometer. 1 s of far-red light (intensity setting 20) was administered on dark-adapted WT and *pgr5-1* leaves. The values were normalised to the maximal oxidation values of PC and P700 and maximal reduction value of Fed obtained by the NIRMAX script. Averaged traces from three biological replicates ± SD are shown. The post-illumination re-oxidation kinetics of Fed were fitted to a first-exponential function (cyan and lime green traces).

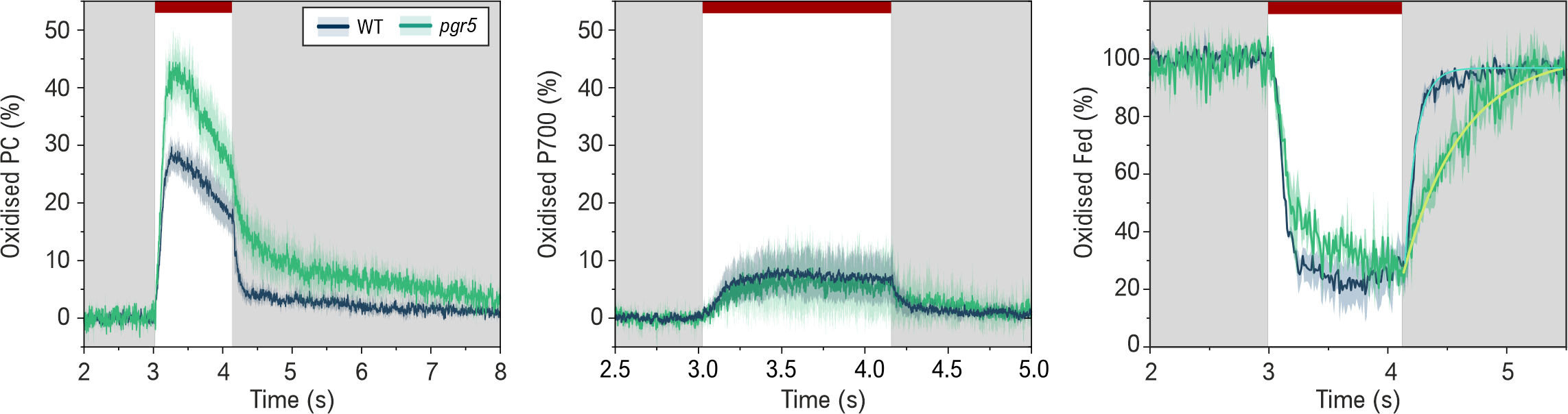

**Supplementary** **Figure S6. *Pmf* dynamics in the *Arabidopsis* *thaliana* WT and *pgr5*-Cas mutant in fluctuating light. (a)** Conductivity of the ATP synthase (gH+), **(b)** magnitude of the *pmf,* and **(c)** proton flux (vH+) in fluctuating light, as determined from dark-interval relaxation kinetics (DIRK) of the ECS signal. ECS was measured from dark-adapted WT-Col-0 and *pgr5-Cas* Arabidopsis leaves under a light regime alternating between 1 min periods of low (50 µmol photons m^-2^s^-1^) and high light (825 µmol photons m^-2^s^-1^). The values are averages from 5 biological replicates ± SEM. The asterisks indicate statistically significant differences between WT and *pgr5*-Cas according to a two-sample Student’s t-tests (P<0.05).

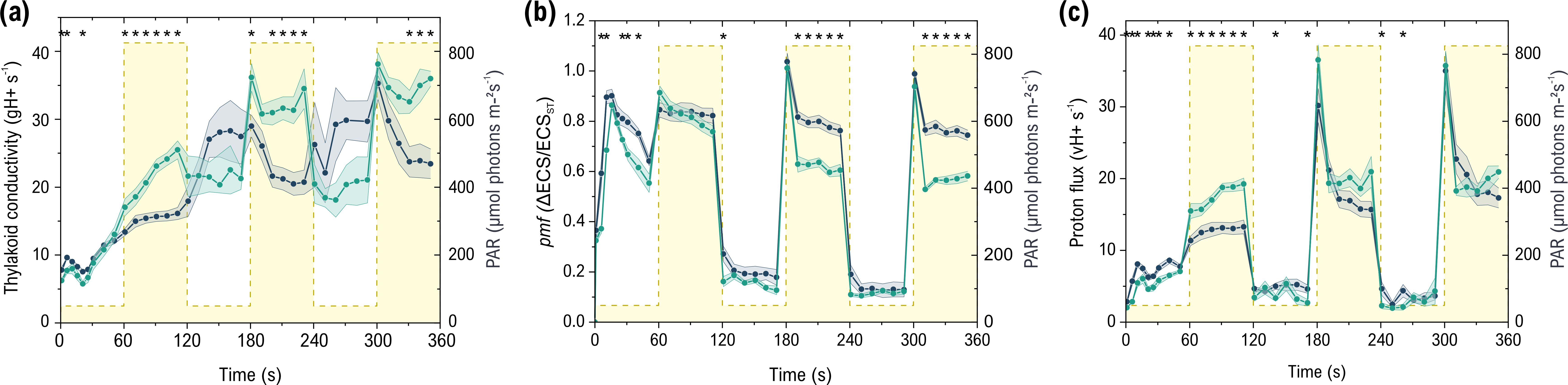

**Supplementary Figure S7. *Pmf* dynamics in the *Arabidopsis* *thaliana* WT and *pgr5*-Cas mutant in fluctuating light in the presence of inhibitors of mitochondrial respiration. (a)** Conductivity of the ATP synthase (gH+), **(b)** magnitude of the *pmf,* and **(c)** proton flux (vH+) in fluctuating light in the presence of 1 mM potassium cyanide (KCN) and 1 mM salicylhydroxamic acid (SHAM), as determined from dark-interval relaxation kinetics (DIRK) of the ECS signal. Detached leaves were placed on water with 1 mM KCN and 1 mM SHAM and incubated in darkness for 2 h. ECS was measured from dark-adapted WT-Col-0 and *pgr5-Cas* Arabidopsis leaves under a light regime alternating between 1 min periods of low (50 µmol photons m^-2^s^-1^) and high light (825 µmol photons m^-2^s^-1^). The values are averages from five biological replicates ± SEM. Statistical significances of the differences between the genotypes at each time point were determined by Student’s T-tests and indicated by asterisks above the traces (P<0.05).

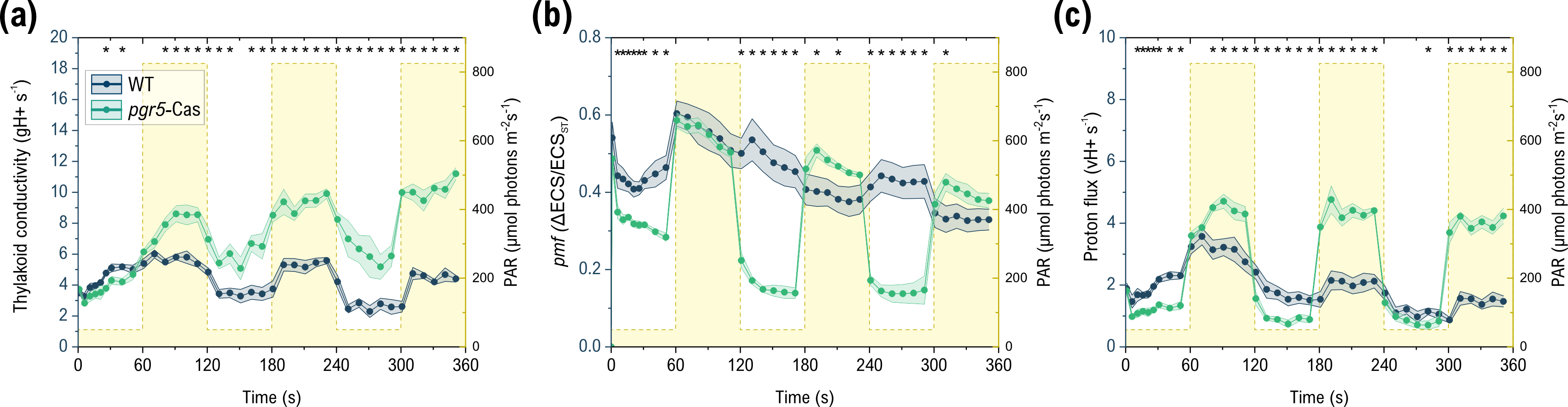

**Supplementary** **Figure S8. Immunodetection of AtpB and Flv3 levels in *Synechocystis* strains.** Two separate clones of the *∆pgr5* and *∆pgr5 flv3* strains were tested. Coomassie brilliant blue staining of the large Rubisco subunit (RbcL) band was used as a loading control. Representative blots from two (AtpB) and one (Flv3) biological replicates are shown. The column graphs show quantification of AtpB (top) and Flv3 (bottom) content in 4 and 2 replicates, respectively (2 or 1 biological and 2 independent clones). Average band intensities normalised to WT intensity +SD are shown with individual replicates as diamonds.

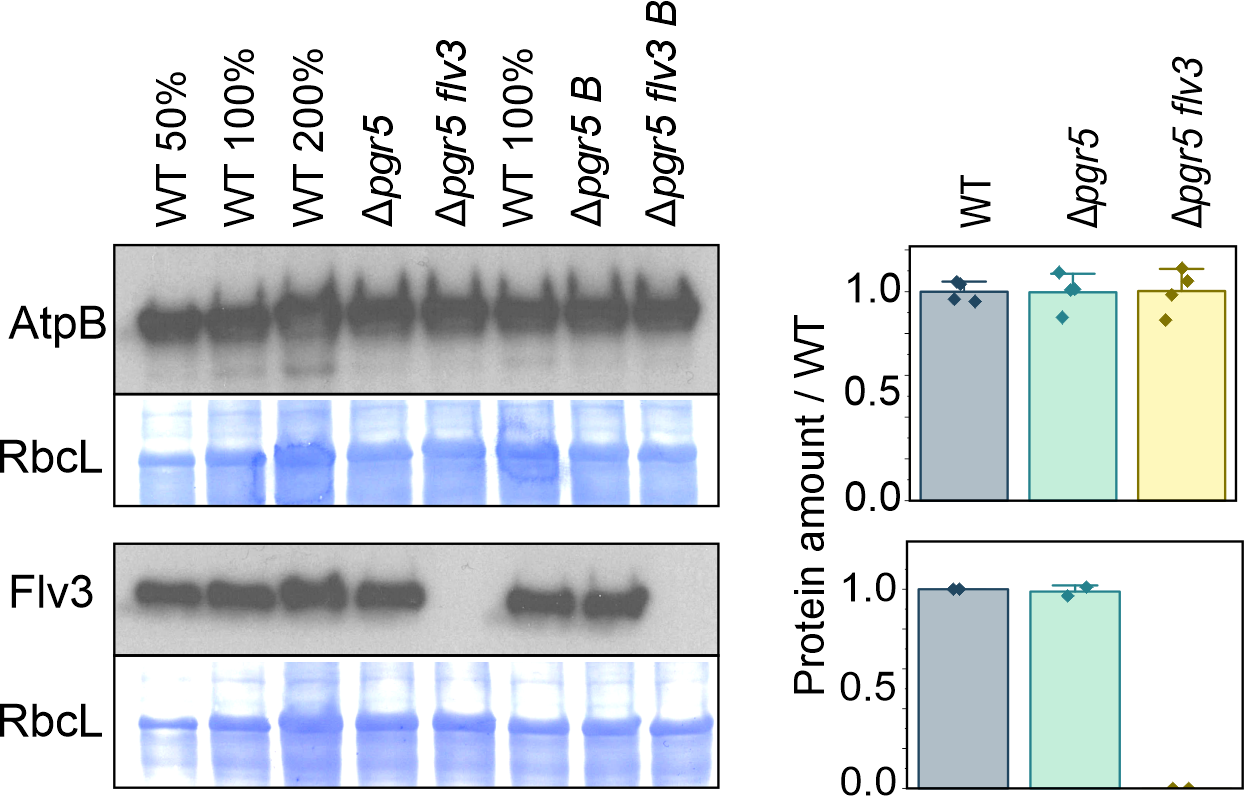

**Supplementary** **Figure S9. (a)** *In vivo* redox kinetics of P700, PC, and Fed in Δ*pgr5 flv3, and* Δ*flv3* strains of *Synechocystis* as monitored with the DUAL-KLAS/NIR spectrophotometer*.* A NIRMAX script adapted for cyanobacteria was used to obtain maximal oxidation of P700 and PC and maximal reduction of Fed. Cells were adjusted to 15 ug Chl/ml and dark-adapted for 10 min prior to the measurements. Representative traces from two biological replicates are shown. **(b)** *In vivo* O_2_ and CO_2_ gas exchange in Δ*pgr5 flv3 and* Δ*flv3 strains of Synechocystis,* as analysed by membrane inlet mass spectrometry (MIMS). Cells were adjusted to 10 µg Chl/ml, and ^18^O_2_ added in equilibrium with ^16^O_2_ in order to distinguish O_2_ uptake from O_2_ evolution. Cells were dark-adapted for 10 min, after which gas exchange was monitored for 5 min in dark, 5 min under 500 µmol photons m^-2^s^-1^ illumination, and 5 min in post-illumination dark. Representative traces from two biological replicates are shown.

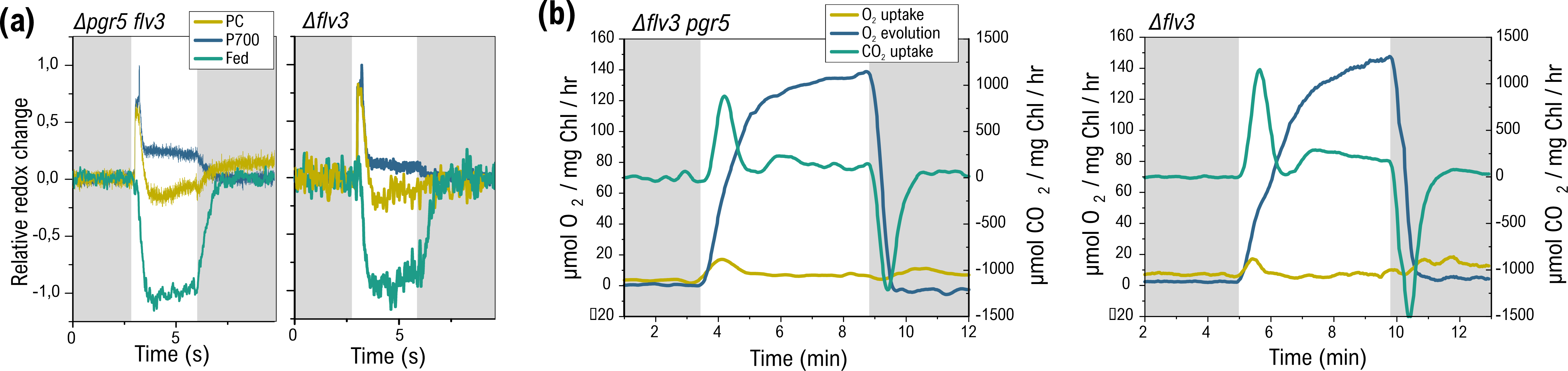

**Supplementary** **Figure S10. P700 oxidation under far-red light.** Deconvoluted redox kinetics of P700 in WT and *Δpgr5* strains of *Synechocystis* as monitored with the DUAL-KLAS/NIR spectrophotometer. Dark-adapted cells were illuminated for 3 s with 3000 µmol photons m^-2^s^-1^, followed by 3 s of darkness and 10 s of illumination under actinic far-red light (FR, intensity 20 in the KLAS-NIR software), of which the one second before to 3 s after the onset of FR are shown. The traces are averages of 7 biological replicates ±SD.

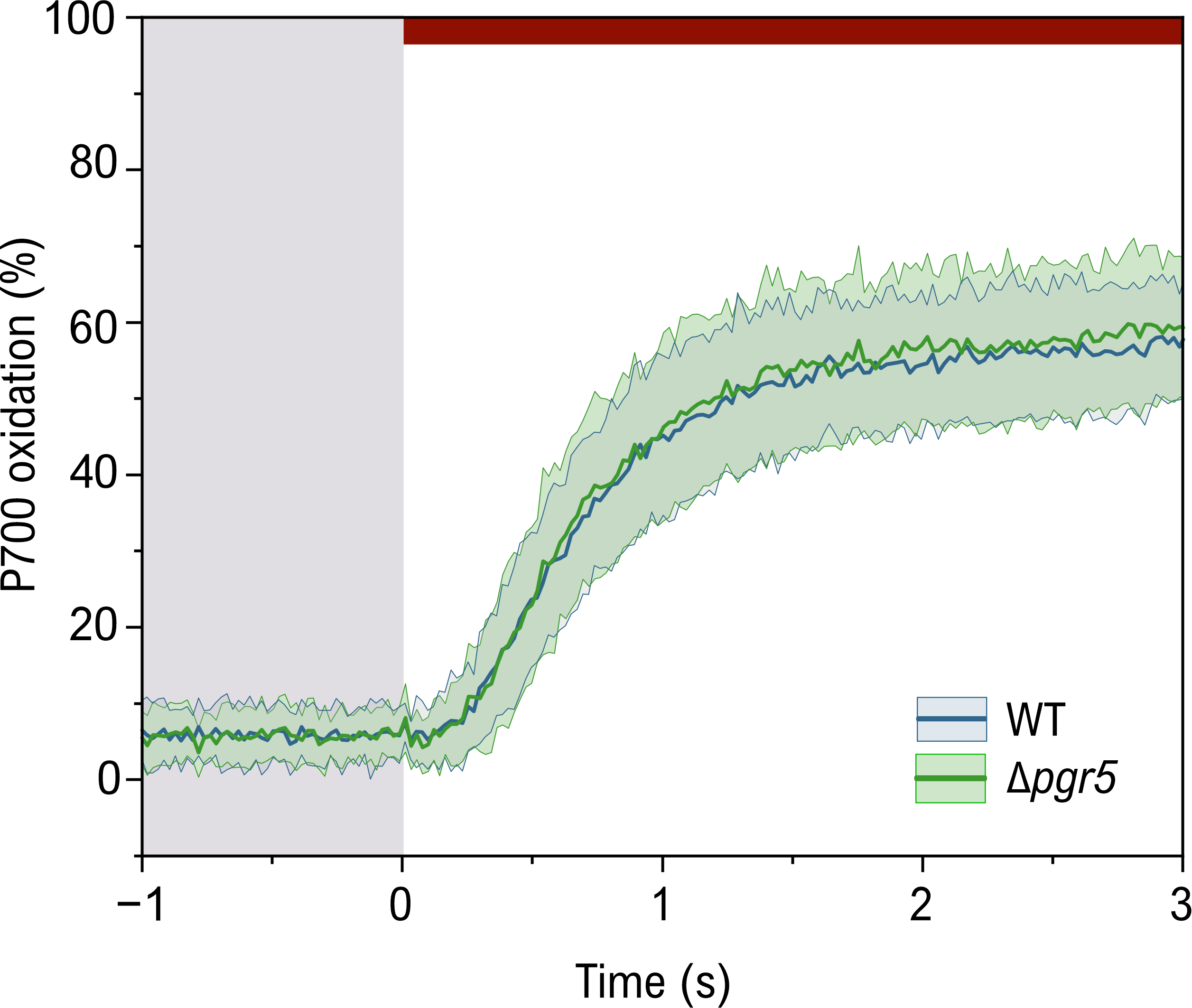

**Supplementary Figure S11. Photosynthetic phenotype of the *Synechocystis Δpgr5* strain. (a)** *In vivo* redox kinetics of P700, PC, and Fed in WT and *Δpgr5* strains of *Synechocystis* as monitored with the DUAL-KLAS/NIR spectrophotometer. Dark-adapted cells were illuminated for 3 s with 3000 µmol photons m^-2^s^-1^. The traces are averages of 7 biological replicates ± SD. **(b)** *In vivo* O_2_ gas exchange in WT and Δ*pgr5* cells, as analysed by membrane inlet mass spectrometry (MIMS). Cells were adjusted to 10 µg Chl/ml, and ^18^O_2_ added in equilibrium with ^16^O_2_ in order to distinguish O_2_ uptake from O_2_ evolution. Cells were dark-adapted for 10 min, after which gas exchange was monitored for 5 min in dark, 5 min under 500 µmol photons m^-2^s^-1^ illumination, and 5 min in post-illumination dark. Averaged traces ±SD from five biological replicates are shown **(c)** Dark respiration rates as averages of O_2_ uptake during the first two minutes of darkness in (b) ± SD (five biological replicates). **(d)** Maximum light-induced O_2_ uptake rates as averages of O_2_ uptake peaks around 30 s of illumination - dark respiration rate ± SD (five biological replicates). **(e)** CO_2_ fixation rates as averages of CO_2_ uptake rates during the final minute in light in (e) + respiration rate during the first two minutes of darkness. Averages +SD from three biological replicates are shown.

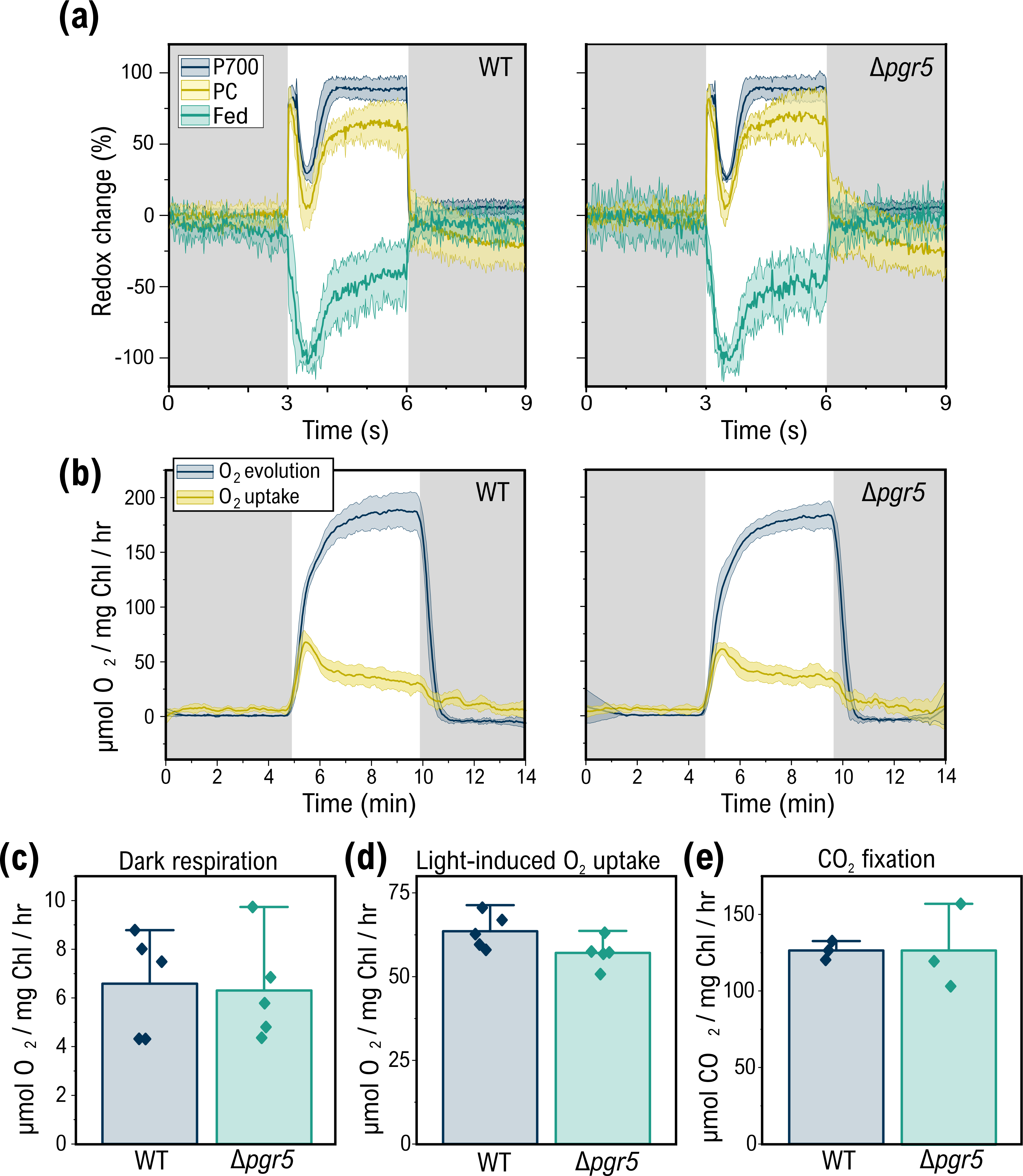

**Supplementary** **Figure S12. *Pmf* dynamics in *Synechococcus elongatus* sp. PCC 7942 under fluctuating light. (A)** Thylakoid conductivity (gH+). **(B)** *pmf* (ΔECS). **(C)** Proton flux (vH+). ECS signal was measured from WT cells as described in Figure 1 legend. Values are means ± SEM from five biological replicates.

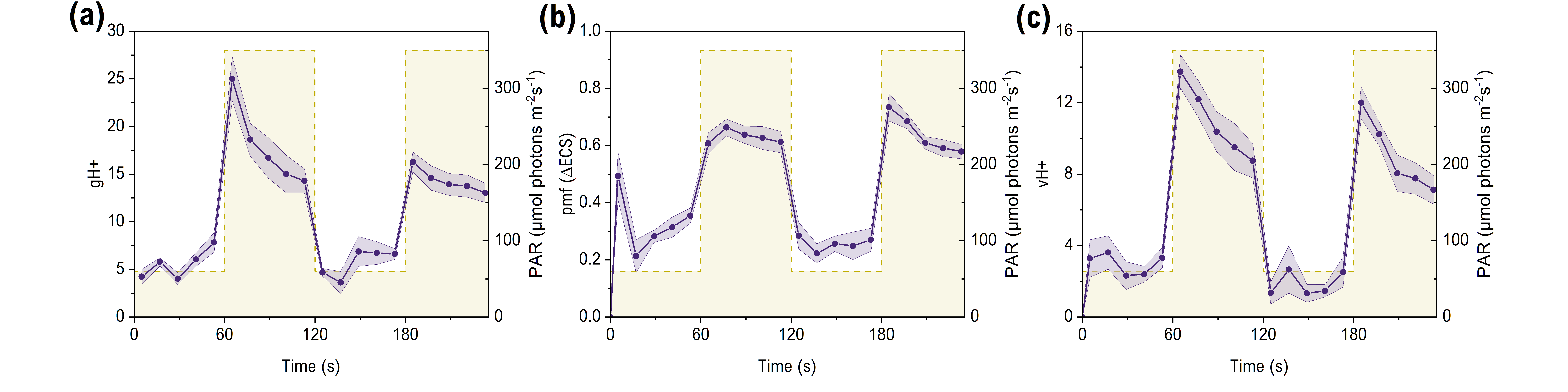

**Supplementary Figure S13. *Pmf* and vH+ dynamics in *Chlamydomonas reinhardtii* under fluctuating light without ECS_ST_ normalisation. (a)** *pmf* (ΔECS). **(b)** Proton flux (vH+). ECS DIRK were measured from WT and *pgr5* knockout cultures under a fluctuating light regime. The values shown are means of three biological replicates ± SEM. Statistical significances of the differences between the genotypes at each time point were determined by Student’s T-tests and indicated by asterisks (P<0.05). The data is derived from Figure 3 and presented without normalising to single-turnover flash-induced ECS magnitude.

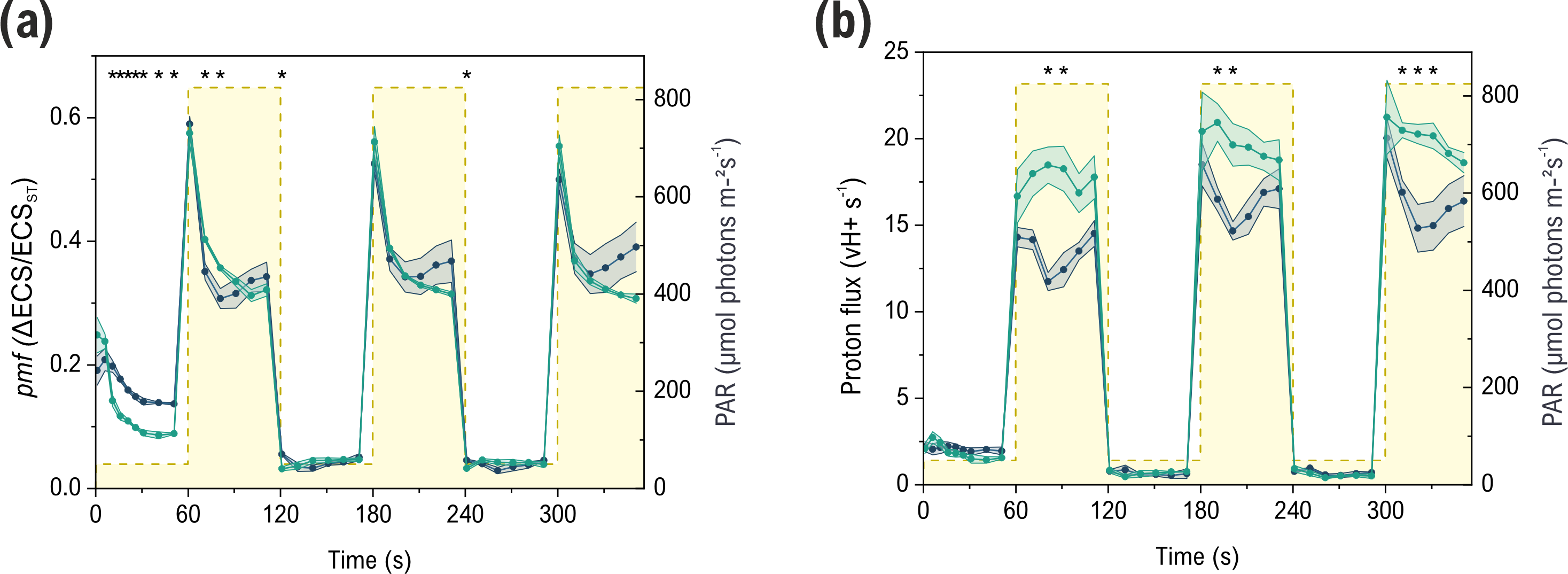

**Supplementary** **Figure S14. *Pmf* dynamics in the C4 model grass *Setaria viridis*. (a)** Thylakoid conductivity (gH+). **(b)** *pmf* (ΔECS). **(c)** Proton flux (vH+). ECS DIRK were measured from WT and *pgr5* knockout plants under a fluctuating light regime. The values shown are means of nine biological replicates ± SEM. Statistical significances of the differences between the genotypes at each time point were determined by Student’s T-tests and indicated by asterisks (P<0.05).

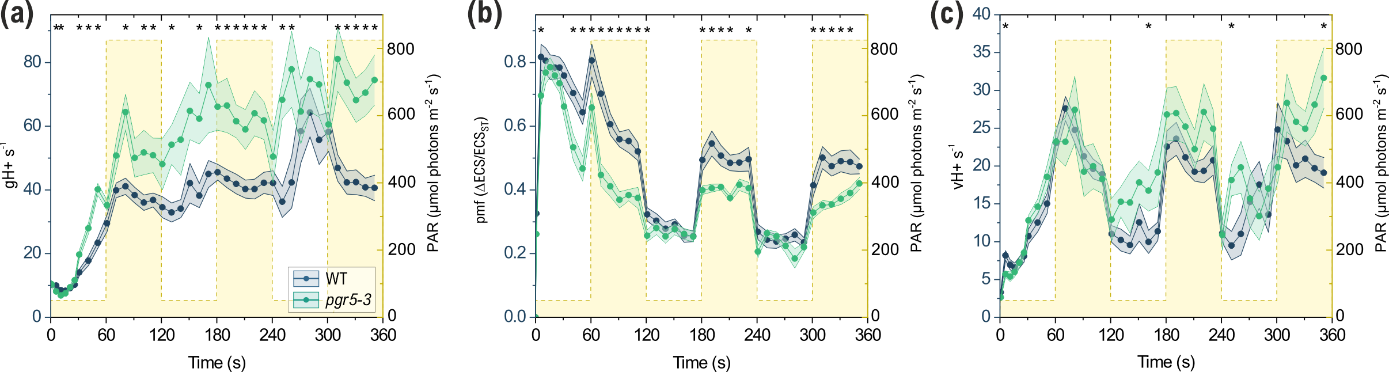

**Supplementary** **Figure S15. Effect of methyl viologen on thylakoid conductivity in *Synechocystis*.** ECS DIRK were measured 5 seconds after the onset of illumination at 500 µmol photons m^-2^ s^-1^. 0.5 mM methyl viologen (MV) was added to the samples in dark prior to the measurements. The values are averages of 4–11 biological replicates +SD.

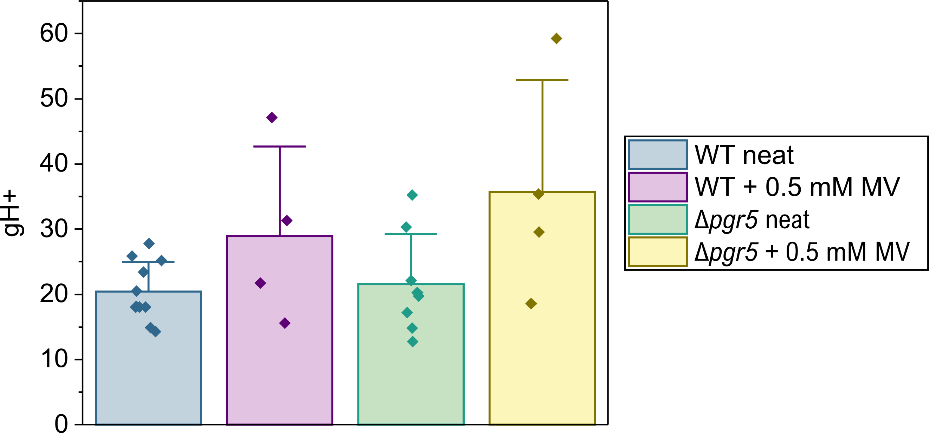

**Supplementary Figure S16. Effect of N-ethylmaleimide (NEM) on thylakoid conductivity under fluctuating light in Arabidopsis**. Extracted from data in Fig. 6c: gH+ at 3 time points during the first, second, and third high light phase was measured in Arabidopsis WT and *pgr5* mutant leaves incubated on water (neat) or water + 0.1 mM NEM. The neat samples are the same as in Figure 1, while the +NEM values are averages of 4 biological replicates +SD with individual replicates shown as diamonds. Statistical significance of differences was tested by one-way ANOVA and Tukey’s post-hoc test of differences of means.

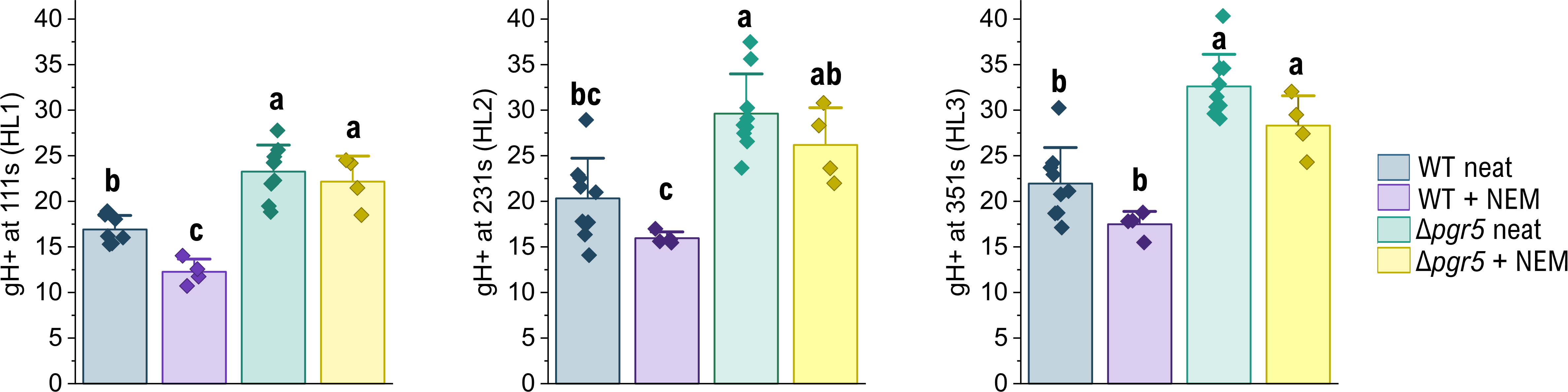

**Supplementary Figure S17. Negative controls for bimolecular fluorescence complementation (BiFC) tests. (a)** BiFC test for protein–protein interaction between Arabidopsis S-subunit of the NDH complex fused to an N-terminal YFP fragment (AtNdhS:YFP-N) and AtPGR5:YFP-C. The left panel shows Chlorophyll *a* autofluorescence in purple, and the right panel YFP fluorescence in yellow. **(b)** BiFC test for protein–protein interaction between AtPGR5:YFP-N and Arabidopsis Thioredoxin x fused to a C-terminal YFP-fragment (AtTRXx:YFP-C).

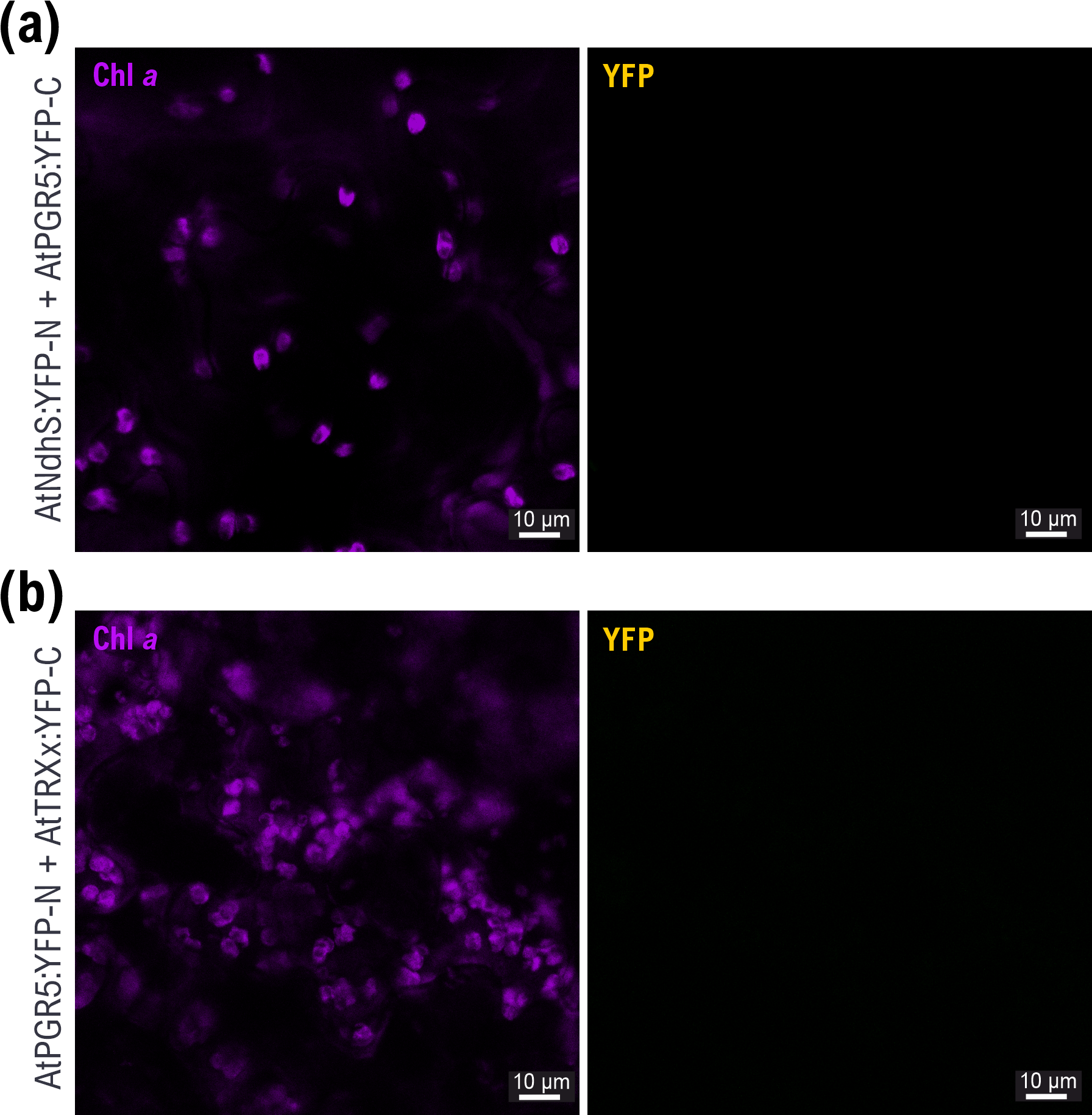

**Supplementary Figure S18. Positive controls for bimolecular fluorescence complementation (BiFC) tests. (a)** AtCF_1_γ:YFP-N + AtTRXf1:YFP-C. The left panel shows Chlorophyll a autofluorescence in purple, the middle panel YFP fluorescence in yellow, and the right panel a merged image of YFP and chlorophyll images. The scale bar is 10 µm. **(b)** AtPGRL1a:YFP-N + AtPGR5:YFP-C. **(c)** Immunoblot against the N-terminal domain of YFP. 20 µg of protein from total protein extracts from uninfected (ctrl) *N. benthamiana* plants, as well as from plants transiently expressing the AtCF_1_γ:YFP-N and AtPGR5:YFP-C fusion proteins (BiFC 1-3). Expected molecular weight of the AtCF_1_γ:YFP-N fusion protein is c.a. 56 kD, while the N-terminal YFP fragment is 17 kD.

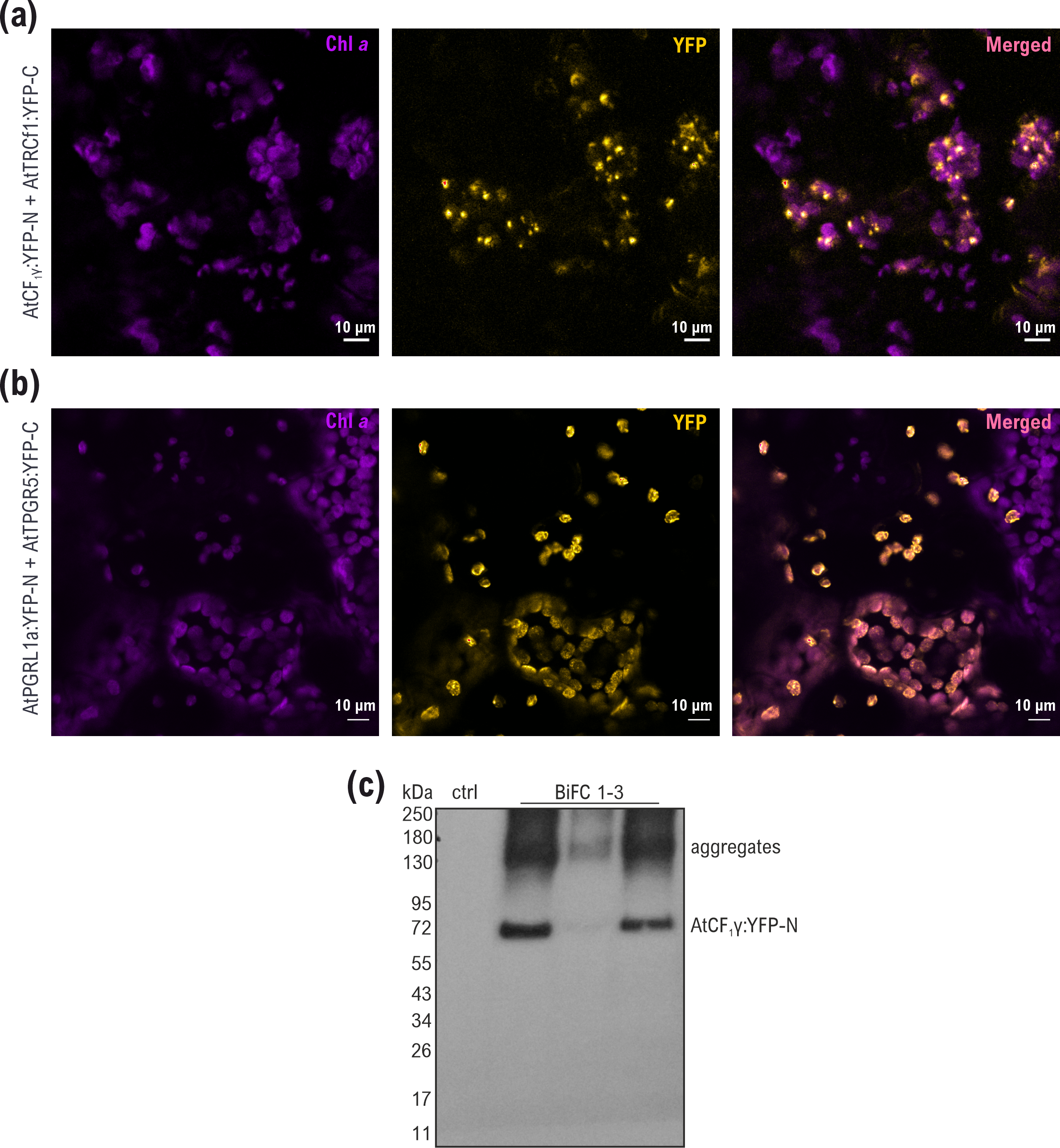

**Table S1. LMEM report table for Arabidopsis WT vs *pgr5-1* (full data)*.*** Linear mixed effects model fitting of gH+ data (presented in Fig. 1) was performed using OriginPro 2025 and the LMEM app extension. LCL=lower confidence limit, UCL=upper confidence limit.

| **Fixed Effect Parameters**  **(Dependent: gH+; continuous: *pmf*; categorical: genotype, light intensity)** | | | | | | |
| --- | --- | --- | --- | --- | --- | --- |
| Intercept | 22.7815 | 1.44595 | 15.7554 | 0.00000 | 19.9425 | 25.6204 |
| *pmf* | -5.91764 | 0.899299 | -6.58028 | 9.27247E-11 | -7.68332 | -4.15196 |
| genotype=*pgr5* | -0.752165 | 1.80192 | -0.417425 | 0.676497 | -4.29004 | 2.78571 |
| PAR=825 | 6.13001 | 0.648026 | 9.45951 | 0.00000 | 4.85768 | 7.40235 |

| **Random Effects (replicate) Variance** | | |
| --- | --- | --- |
|  | **Variance** | **Standard Deviation** |
| Intercept | 12.8296 | 3.58185 |
| Residuals | 58.8768 | 7.67312 |

| **Statistics** | |
| --- | --- |
|  | **gH+** |
| Number of Points | 697.000 |
| Degrees of Freedom | 692.000 |
| Reduced Chi-Sqr | 57.7464 |
| Residual Sum of Squares | 39960.5 |
| R Value | 0.518127 |
| Adj. R-Square | 0.264227 |
| Intraclass Correlation | 0.178919 |

**Table S2. LMEM report table for Arabidopsis WT vs *pgr5-1* (high light data)*.*** Linear mixed effects model fitting of gH+ data (presented in Fig. 1) was performed using OriginPro 2025 and the LMEM app extension. LCL=lower confidence limit, UCL=upper confidence limit.

| **Fixed Effect Parameters**  **(Dependent: gH+; continuous: *pmf*; categorical: genotype)** | | | | | | |
| --- | --- | --- | --- | --- | --- | --- |
|  | **Value** | **Standard Error** | **t-Value** | **Prob>\|t\|** | **95% LCL** | **95% UCL** |
| Intercept | 20.018 | 1.747 | 11.455 | 0.000 | 16.580 | 23.455 |
| *pmf* | 0.406 | 1.196 | 0.339 | 0.735 | -1.947 | 2.759 |
| genotype=*pgr5* | 5.620 | 1.736 | 3.237 | **0.001** | 2.205 | 9.035 |

| **Random Effects (replicate) Variance** | | |
| --- | --- | --- |
|  | **Variance** | **Standard Deviation** |
| Intercept | 11.67 | 3.42 |
| Residuals | 23.13 | 4.81 |

| **Statistics** | |
| --- | --- |
|  | **gH+** |
| Number of Points | 337.00 |
| Degrees of Freedom | 333.00 |
| Reduced Chi-Sqr | 22.20 |
| Residual Sum of Squares | 7391.72 |
| R Value | 0.68 |
| Adj. R-Square | 0.45 |
| Intraclass Correlation | 0.34 |

**Table S3. LMEM report table for Arabidopsis WT vs *pgr5-*Cas (full data)*.*** Linear mixed effects model fitting of gH+ data (presented in Fig. 1) was performed using OriginPro 2025 and the LMEM app extension. LCL=lower confidence limit, UCL=upper confidence limit.

| **Fixed Effect Parameters**  **(Dependent: gH+; continuous: *pmf*; categorical: genotype, light intensity)** | | | | | | |
| --- | --- | --- | --- | --- | --- | --- |
|  | **Value** | **Standard Error** | **t-Value** | **Prob>\|t\|** | **95% LCL** | **95% UCL** |
| Intercept | 25.4671 | 1.39064 | 18.3132 | 0.00000 | 22.7325 | 28.2017 |
| *pmf* | -22.2754 | 1.55469 | -14.3279 | 0.00000 | -25.3326 | -19.2182 |
| genotype=*pgr5* | 0.0229560 | 1.64618 | 0.0139450 | 0.988881 | -3.21415 | 3.26006 |
| PAR=825 | 17.3324 | 0.902738 | 19.1998 | 0.00000 | 15.5572 | 19.1076 |

| **Random Effects (replicate) Variance** | | |
| --- | --- | --- |
|  | **Variance** | **Standard Deviation** |
| Intercept | 5.50523 | 2.34632 |
| Residuals | 45.4273 | 6.73998 |

| **Statistics** | |
| --- | --- |
|  | **gH+** |
| Number of Points | 373.000 |
| Degrees of Freedom | 368.000 |
| Reduced Chi-Sqr | 44.7426 |
| Residual Sum of Squares | 16465.3 |
| R Value | 0.738860 |
| Adj. R-Square | 0.540978 |
| Intraclass Correlation | 0.108089 |

**Table S4. LMEM report table for Arabidopsis WT vs *pgr5-*Cas (high light only)*.*** Linear mixed effects model fitting of gH+ data (presented in Fig. 1) was performed using OriginPro 2025 and the LMEM app extension. LCL=lower confidence limit, UCL=upper confidence limit.

| **Fixed Effect Parameters**  **(Dependent: gH+; continuous: *pmf*; categorical: genotype)** | | | | | | |
| --- | --- | --- | --- | --- | --- | --- |
|  | **Value** | **Standard Error** | **t-Value** | **Prob>\|t\|** | **95% LCL** | **95% UCL** |
| Intercept | 28.8582 | 3.58372 | 8.05257 | 1.17462E-13 | 21.7856 | 35.9308 |
| *pmf* | -8.56183 | 3.99167 | -2.14493 | 0.0333301 | -16.4395 | -0.684135 |
| genotype=*pgr5* | 6.89413 | 2.09983 | 3.28319 | **0.00123811** | 2.75005 | 11.0382 |

| **Random Effects (replicate) Variance** | | |
| --- | --- | --- |
|  | **Variance** | **Standard Deviation** |
| Intercept | 8.19908 | 2.86340 |
| Residuals | 42.2560 | 6.50046 |

| **Statistics** | |
| --- | --- |
|  | **gH+** |
| Number of Points | 180.000 |
| Degrees of Freedom | 176.000 |
| Reduced Chi-Sqr | 41.0078 |
| Residual Sum of Squares | 7217.38 |
| R Value | 0.618942 |
| Adj. R-Square | 0.372573 |
| Intraclass Correlation | 0.162503 |

**Table S5. LMEM report table for *Chlamydomonas* WT vs *pgr5* (full data)*.*** Linear mixed effects model fitting of gH+ data (presented in Supplementary Figure S14) was performed using OriginPro 2025 and the LMEM app extension. LCL=lower confidence limit, UCL=upper confidence limit.

| **Fixed Effect Parameters**  **(Dependent: gH+; continuous: *pmf*; categorical: genotype, light intensity)** | | | | | | |
| --- | --- | --- | --- | --- | --- | --- |
|  | **Value** | **Standard Error** | **t-Value** | **Prob>\|t\|** | **95% LCL** | **95% UCL** |
| Intercept | 8.64301 | 1.57026 | 5.50417 | 1.00303E-7 | 5.54878 | 11.7372 |
| *pmf* | 11.9195 | 1.37637 | 8.66008 | 8.88178E-16 | 9.20732 | 14.6316 |
| genotype=*pgr5* | 5.71666 | 1.90817 | 2.99588 | **0.00304169** | 1.95658 | 9.47675 |
| PAR=825 | 21.7050 | 1.40815 | 15.4139 | 0.00000 | 18.9302 | 24.4798 |

| **Random Effects (replicate) Variance** | | |
| --- | --- | --- |
|  | **Variance** | **Standard Deviation** |
| Intercept | 3.46698 | 1.86198 |
| Residuals | 76.7083 | 8.75833 |

| **Statistics** | |
| --- | --- |
|  | **gH+** |
| Number of Points | 231.000 |
| Degrees of Freedom | 226.000 |
| Reduced Chi-Sqr | 76.1865 |
| Residual Sum of Squares | 17218.2 |
| R Value | 0.873191 |
| Adj. R-Square | 0.758259 |
| Intraclass Correlation | 0.0432425 |

**Table S6. LMEM report table for *Setaria viridis* WT vs *pgr5-1* (full data)*.*** Linear mixed effects model fitting of gH+ data (presented in Supplementary Figure S14) was performed using OriginPro 2025 and the LMEM app extension. LCL=lower confidence limit, UCL=upper confidence limit.

| **Fixed Effect Parameters**  **(Dependent: gH+; continuous: *pmf*; categorical: genotype, light intensity)** | | | | | | |
| --- | --- | --- | --- | --- | --- | --- |
|  | **Value** | **Standard Error** | **t-Value** | **Prob>\|t\|** | **95% LCL** | **95% UCL** |
| Intercept | 61.1700 | 3.92034 | 15.6033 | 0.00000 | 53.4729 | 68.8671 |
| *pmf* | -67.5507 | 3.44556 | -19.6051 | 0.00000 | -74.3156 | -60.7857 |
| genotype=*pgr5* | 10.6642 | 4.13429 | 2.57946 | **0.0100996** | 2.54705 | 18.7814 |
| PAR=825 | 17.2862 | 4.12493 | 4.19066 | 3.13973E-5 | 9.18738 | 25.3850 |

| **Random Effects (replicate) Variance** | | |
| --- | --- | --- |
|  | **Variance** | **Standard Deviation** |
| Intercept | 137.655 | 11.7327 |
| Residuals | 294.045 | 17.1477 |

| **Statistics** | |
| --- | --- |
|  | **gH+** |
| Number of Points | 701.000 |
| Degrees of Freedom | 696.000 |
| Reduced Chi-Sqr | 281.926 |
| Residual Sum of Squares | 196220 |
| R Value | 0.768850 |
| Adj. R-Square | 0.588781 |
| Intraclass Correlation | 0.318867 |

**Table S7. LMEM report table for *Synechocystis* WT vs Δ*pgr5* (full data)*.*** Linear mixed effects model fitting of gH+ data (presented in Supplementary Figure S14) was performed using OriginPro 2025 and the LMEM app extension. LCL=lower confidence limit, UCL=upper confidence limit.

| **Fixed Effect Parameters**  **(Dependent: gH+; continuous: *pmf*; categorical: genotype, light intensity)** | | | | | | |
| --- | --- | --- | --- | --- | --- | --- |
|  | **Value** | **Standard Error** | **t-Value** | **Prob>\|t\|** | **95% LCL** | **95% UCL** |
| Intercept | 3.63450 | 3.35216 | 1.08423 | 0.280367 | -3.00036 | 10.2694 |
| *pmf* | -1.31544 | 3.52226 | -0.373465 | 0.709440 | -8.28699 | 5.65610 |
| genotype=Δ*pgr5* | 12.3331 | 5.01787 | 2.45783 | **0.0153594** | 2.40133 | 22.2649 |
| PAR=350 | 13.4853 | 1.58001 | 8.53490 | 4.17444E-14 | 10.3580 | 16.6126 |

| **Random Effects (replicate) Variance** | | |
| --- | --- | --- |
|  | **Variance** | **Standard Deviation** |
| Intercept | 43.4914 | 6.59480 |
| Residuals | 53.3637 | 7.30505 |

| **Statistics** | |
| --- | --- |
|  | **gH+** |
| Number of Points | 129.000 |
| Degrees of Freedom | 124.000 |
| Reduced Chi-Sqr | 51.4092 |
| Residual Sum of Squares | 6374.74 |
| R Value | 0.829096 |
| Adj. R-Square | 0.677317 |
| Intraclass Correlation | 0.449035 |

**Table S8. Hydrogen Bond Interactions and salt bridges between AtPGR5 and SoCF_1_γ in the ClusPro 2.0 docking model.** Interaction interface was examined using the PDBePISA online tool.

**Hydrogen bonds**

| AtPGR5  residue | Atom | SoCF_1_γ  residue | Atom | Distance (Å) |
| --- | --- | --- | --- | --- |
| LYS 92 | HZ2 | **ASN 81** | OD1 | 1.66984 |
| LYS 92 | HZ3 | **GLU 77** | OE1 | 1.80101 |
| LYS 92 | HZ1 | **GLU 77** | OE2 | 1.75972 |
| ALA 93 | O | **ARG 74** | HH22 | 1.77652 |
| ALA 93 | O | **ARG 74** | HE | 1.92399 |
| GLN 99 | OE1 | **ARG 73** | HH11 | 1.79092 |
| PHE 131 | O | **LYS 71** | HZ1 | 1.71906 |

**Salt bridges**

| AtPGR5  residue | Atom | SoCF_1_γ  residue | Atom | Distance (Å) |
| --- | --- | --- | --- | --- |
| LYS 92 | NZ | **GLU 77** | OE1 | 2.59516 |
| LYS 92 | NZ | **GLU 77** | OE2 | 2.60983 |

**Table S9. Hydrogen Bond Interactions between SynCF_1_γ and SynPgr5 in the AlphaFold2 model.** Interaction interface was examined using the PDBePISA online tool**.**

| SynCF_1_γ | Atom | SynPgr5 | Atom | Distance (Å) |
| --- | --- | --- | --- | --- |
| ARG 10 | NH1 | **ALA 3** | O | 3.40 |
| ASN 16 | ND2 | **ASN 49** | O | 2.88 |
| VAL 14 | N | **LEU 53** | O | 2.92 |
| VAL 14 | N | **SER 56** | OG | 3.32 |
| ARG 10 | NE | **ASN 57** | O | 3.75 |
| ARG 10 | NH2 | **ASN 57** | O | 3.62 |
| PRO 2 | O | **ARG 10** | NH2 | 3.74 |
| ASN 3 | O | **ARG 10** | NH2 | 2.60 |
| ALA 6 | O | **ARG 22** | NH2 | 2.02 |
| ARG 10 | O | **ASN 57** | N | 3.57 |
| SER 13 | O | **ALA 54** | N | 3.45 |
| SER 13 | OG | **ALA 54** | N | 2.77 |
| THR 17 | OG1 | **ALA 54** | N | 2.66 |

**Table S10. Alignment of the ATP synthase F_1_γ amino acid sequences from *Synechocystis* sp. PCC 6803 and *Synechococcus elongatus* sp. PCC 7942.** The alignment was performed using the NCBI protein blast tool. The residues highlighted with yellow are predicted to be involved in forming hydrogen bonds with SynPGR5, while the alanine residue marked with magenta constitutes a different residue in the Synechococcus sequence.

```
Synechocystis  1    MPNLKAIRDRIQSVKNTKKITEAMRLVAAAKVRRAQEQVLSTRPFADALAQVLYNLQNRL  60
Synechococcus  1    MANLKAIRDRIKSVRNTRKITEAMRLVAAAKVRRAQEQVLSTRPFADRLAQVLAGLQQRL  60

Synechocystis  61   SFAETELPLFEQREPKAVALLVVTGDRGLCGGYNVNAIKRAEQRAKELKNQGIAVKLVLV  120
Synechococcus  61   QFENVDLPLLQRREVKTVALLVVSGDRGLCGGYNSNVIRRAEQRARELSAQGLDYKFVIV  120

Synechocystis  121  GSKAKQYFGRRDYDVAASYANLEQIPNASEAAQIADSLVALFVSETVDRVELIYTRFVSL  180
Synechococcus  121  GRKAGQYFQRREQPIEATYSGLEQIPTAQEANDIADELLSLFLSGTVDRVELVYTKFLSL  180

Synechocystis  181  ISSQPVVQTLFPLSPQGLEAPDDEIFRLITRGGKFQVEREKVEAPVESFPQDMIFEQDPV  240
Synechococcus  181  VASNPVVQTLLPLDPQGLASSDDEIFRLTTRGGSFTVEREKLTSEVAPLPRDMIFEQDPA  240

Synechocystis  241  QILEALLPLYNTNQLLRALQESAASELAARMTAMSNASDNAGQLIGTLTLSYNKARQAAI  300
Synechococcus  241  QILSALLPLYLSNQLLRALQEAAASELAARMTAMNSASDNANALVGQLTLVYNKARQAAI  300

Synechocystis  301  TQELLEVVAGANSL  314
Synechococcus  301  TQELLEVVAGAEAL  314
```
